# Neuromesodermal progenitor origin of trunk neural crest *in vivo*

**DOI:** 10.1101/2021.02.10.430513

**Authors:** Martyna Lukoseviciute, Sarah Mayes, Tatjana Sauka-Spengler

**Affiliations:** University of Oxford, MRC Weatherall Institute for Molecular Medicine, Radcliffe Department of Medicine, Oxford OX3 9DS, UK; Department of Cell and Molecular Biology, Karolinska Institutet, SE-171 77 Stockholm, Sweden

**Keywords:** neural crest, neuromesodermal progenitors, gene regulatory network, axial progenitors, embryo patterning

## Abstract

Neural crest (NC) is a vertebrate-specific population of multipotent embryonic cells predisposed to diverse derivatives along the anteroposterior (A-P) axis. Only cranial NC progenitors give rise to ectomesenchymal cell types, whereas trunk NC is biased for neuronal cell fates. By integrating multimodal single-cell analysis, we provide evidence for divergent embryonic origins of cranial vs. trunk NC that explain this dichotomy. We show that the NC regulator foxd3 is heterogeneously expressed across the A-P axis and identify its specific cranial and trunk autoregulatory enhancers. Whereas cranial-specific enhancer is active in the *bona fide* NC, the trunk foxd3 autoregulatory element surprisingly marked bipotent tailbud neuromesodermal progenitors (NMps). We integrated NMp single cell epigemomics and trasncriptomics data and for the first time reconstructed anamniote NMp gene regulatory network. Moreover, using pseudotime and developmental trajectory analyses of NMps and NC during normal development and in *foxd3* mutants, we demonstrate an active role for foxd3 in balancing non-cranial NC and NMp fates during early embryonic development. Strikingly, we show that a portion of posterior NC in the developing zebrafish embryo is derived from the pro-neural NMps. This suggests a common embryonic origin of trunk NC and NM progenitors that is distinct from cranial NC anlage, and elucidates pro-neural bias of trunk NC.

## Introduction

The neural crest (NC) is a unique multipotent cell population that gives rise to an extraordinary diversity of tissues in vertebrate embryos, including chondrocytes and osteocytes of the craniofacial skeleton, neurons and glia of the peripheral nervous system, and all skin pigment cells in the body.

NC population originates at the end of gastrulation when three archetypal germ layers, ectoderm, mesoderm and endoderm, are definitively formed. Due to relatively late specification during embryogenesis and a unique capacity to generate a prodigious number of derivatives including mesenchymal, neuronal and secretory cell types, NC is sometimes considered a 4th germ layer. NC is thought to arise uniformly from a narrow stretch of ectoderm termed neural plate border, situated between the future neural and non-neural ectoderm. As neurulation proceeds, neural plate border progenitors come to reside within the dorsal aspect of the newly formed neural tube, from which they delaminate, migrate to precise destinations, and differentiate into distinct derivatives. As the NC cells are generated in the entire embryo, their location along the anterior-posterior (A-P) axis influences their fate (Le Douarin, 1999). For instance, the cranial NC preference for mesenchymal fates results in facial cartilage and bone formation, whereas trunk NC displays a bias for neuronal fates, such as sensory neurons of the dorsal root ganglia (Etchevers et al., 2019; Le Douarin, 1999). Some distinct transcription factors that are capable of driving cranial NC identity in the trunk (e.g. *Sox8*, *Tfap2b*, and *Ets1*) have been described (Simões-Costa and Bronner, 2016) but efforts to re-program trunk NC towards cranial NC were only partially successful, with limited activation of ectomesenchymal properties in the trunk NC (Simões-Costa and Bronner, 2016; Soldatov et al., 2019). Such meticulous differences between cranial NC and trunk NC fate potentials raise a possibility of distinct embryonic origins.

Correspondingly, *in vitro* NC models derived by human embryonic stem cell differentiation yield mostly cranial-like NC and not trunk-like NC (Bajpai et al., 2010). Moreover, human pluripotent cells (hPSCs) have so far been successfully differentiated into trunk-like NC only via a neuromesodermal progenitor (NMp) intermediate (Frith et al., 2018). NMps are bipotent self-renewing cells that reside in the primitive streak region at the posterior of the developing embryo, capable of giving rise to both paraxial mesoderm and spinal cord neural derivatives during amniote axis elongation (Tzouanacou et al., 2009). Additional attempts to model NC development using *in vitro* differentiation of hPSCs, using Wnt and FGF as the main drivers of trunk NC identity, also revealed that *in vitro* trunk-like NC progenitors exhibit a transient intermediate state, expressing both early NC (*Msx1/2*, *Pax3*, *Zic1/3*) and NMp markers (*Bra*, *Sox2*, *Msgn1*) (Frith et al., 2018; Gomez et al., 2019; Hackland et al., 2019). However, such *in vitro* differentiation assays are solely based on user-selected and supplied differentiation cocktails, are taken out of embryonic context and overall represent artificial systems. Thus far, a distinctive trunk NC territory of origin, different from described border region from which cranial NC derive, has not been described *in vivo*. Fate mapping and lineage tracing data from the amniote primitive streak/tailbud, where NMps localise in amniotes, have hinted that tailbud region contributes to trunk NC generation *in vivo* (Anderson et al., 2013; Javali et al., 2017; McGrew et al., 2008; Rodrigo Albors et al., 2018; Tzouanacou et al., 2009; Wymeersch et al., 2016). However, these classical isotopic graft and labelling experiments suggested that lumbosacral NC arise from a heterogenous tailbud region shared with other axial progenitors near the primitive streak (Catala et al., 1995; Schoenwolf and Nichols, 1984) and have not instigated the common origin of trunk NC and the NMps. Hence whether embryonic trunk NC progenitors transit via an NMp intermediate state, undertake an alternative route or originate from the neural plate border as cranial NC do, remains to be resolved.

To address this long-standing question, we investigated NC heterogeneity in anamniote zebrafish embryo. We isolated cells that express a critical NC regulator, foxd3, expressed across the entire anteroposterior (A-P) axis in all vertebrates and have performed and integrated single-cell multimodal analysis to uncover underlying NC heterogeneity at transcriptional, epigenomic and *cis*-regulatory levels. We discover that unlike in avians where anterior and posterior FoxD3 regulation delineates anterior and posterior NC populations, respectively (Simões-Costa et al., 2012), zebrafish exhibit more complex differential regulation of foxd3 along the anterior-posterior axis. While tthe anterior autoregulatory *foxd3* enhancer lit up cranial NC, the reporter for trunk-specific *foxd3* enhancer activity did not label the *bona fide* trunk NC. Instead, the tailbud NMps and early NMp derivatives exhibited trunk-specific *foxd3* enhancer activity. By integrating single-cell transcriptional and epigenomic profiles we reconstructed the core gene regulatory networks (GRNs) underlying NMp differentiation. Finally, we identified early tailbud NMp-derived pro-neural cells that give rise to prospective trunk NC cells. Such transitioning NMp/NC cells express neural plate border and early NC genes in addition to pro-neural NMp genes and are later located within neuronal trunk derivative structures such as dorsal root ganglia. This intermediate pro-neural trunk NMp/NC cell population is decreased in the *foxd3* mutant, leading to the expansion of the naïve tailbud NMps instead, emphasising a conserved foxd3 role in NC specification. Thus, this work, for the first time, shows NMp-neural cell contribution to pro-neuronal trunk NC *in vivo*, shedding light on its distinctive embryonic origin from that of cranial NC and thus explaining the trunk NC bias for neuronal fates.

## Results

### Distinct foxd3 regulatory landscapes in the cranial NC and tail NMps

Foxd3 acts both as a chromatin/transcriptional activator and a repressor in the NC (Krishnakumar et al., 2016; Lukoseviciute et al., 2018; Respuela et al., 2016) and is required for NC induction (Hanna et al., 2002; Teng et al., 2008; Thomas and Erickson, 2009; Tompers et al., 2005). To uncover the heterogeneity of distinct populations of foxd3-expressing cells along the A-P axis during NC development in zebrafish, we FACS-isolated citrine+/foxd3+ cells at all axial levels from neurula stage *Gt(foxd3-citrine)*^*ct110a*^ zebrafish embryos (5-6 somite stage (ss)) (Hochgreb-Hägele and Bronner, 2013).

Single-cell RNA-sequencing (scRNA-seq) followed by filtering and clustering of the sequenced cells (fig. S1, A-C), revealed three major foxd3+ cell types: NC (*tfap2a*, *pax3a*, *sox10*), including cranial NC (*otx1+*, *otx2b+*) and trunk NC (*otx1-*, *otx2b-*), hindbrain NC (*mafba*, *egr2b*, *hoxb3a*) and NMp-related cells (*tbxta*, *sox2*, *msgn1*) (fig. 1 A-B, S1 B). Whereas foxd3 is a well-known NC regulator (Stewart et al., 2006), its role in NMp development has not been documented, hence the presence of foxd3+ NMp-related cells was unexpected.

**Figure 1:**
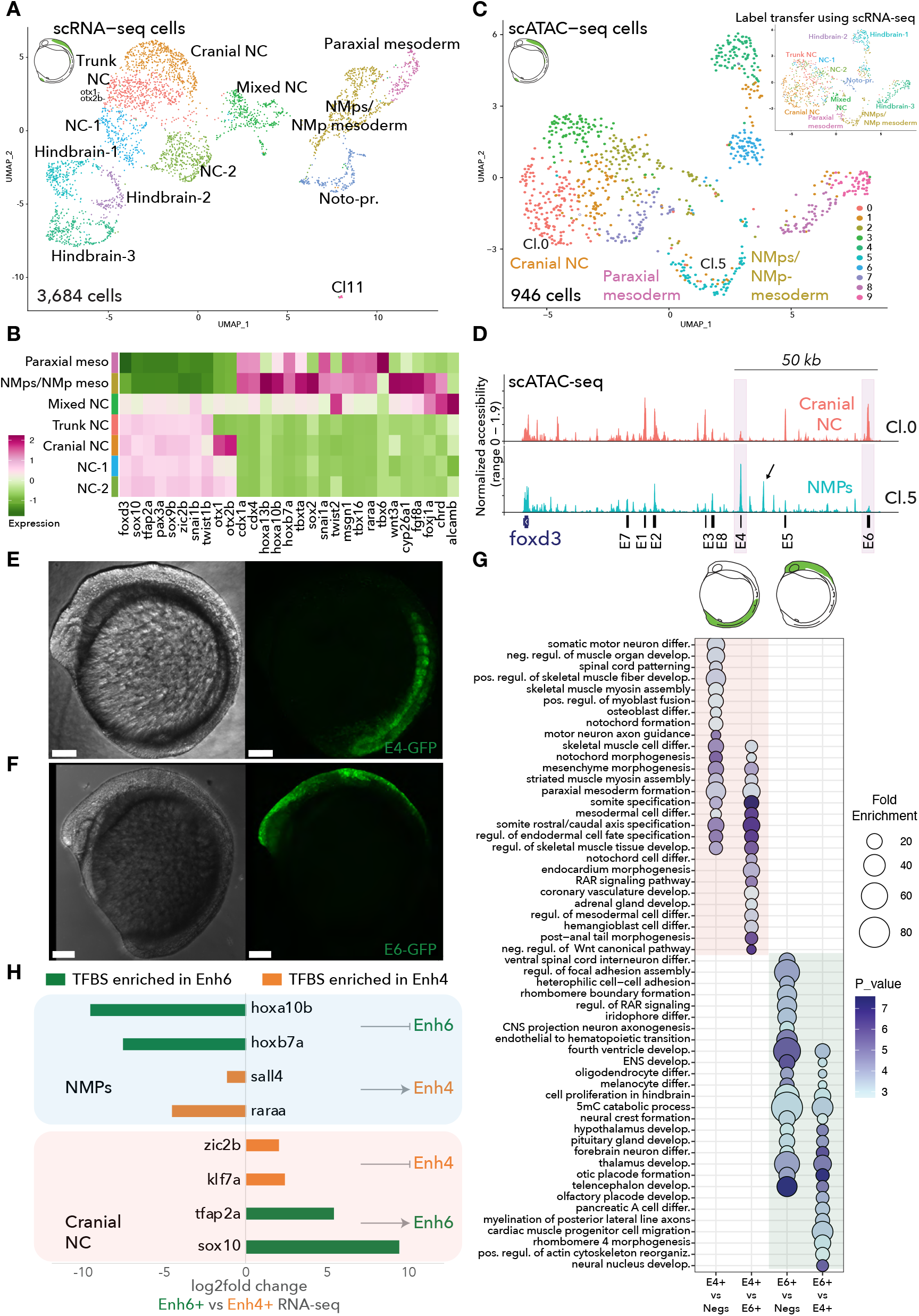
Foxd3+ cell heterogeneity and associated cell chromatin accessibility across the anteroposterior axis in the 5-6 somite stage (ss) zebrafish embryo. (**A**) Uniform Manifold Approximation and Projection (UMAP) embedding showing 12 clusters of foxd3+ 3,684 single cell transcriptomes (scRNA-seq) enriched during fluorescence-activated cell sorting (FACS) from 5-6ss *Gt(foxd3-citrine)*^*ct110a*^ transgenic embryos. Clusters were manually annotated based on the enriched cell-type markers (S1 B). NC – neural crest, NMps – neuromesodermal progenitors, Noto-pr. – notochord progenitors, Cl – cluster. (**B**) Heatmap illustrating selected cluster from (**A**) average expression of NC and NMp associated genes. (**C**) UMAP embedding showing 10 clusters based on single cell chromatin accessibility (scATAC-seq) similarities of foxd3+ 946 cells enriched during FACS from 5-6ss *Gt(foxd3-citrine)*^*ct110a*^ transgenic embryos. Anchors found between scRNA-seq (**A**) and scATAC-seq datasets were used to transfer scRNA-seq cluster labels onto the scATAC-seq clusters, displayed on the top right. (**D**) Chromatin pseudo-bulk accessibility profiles of *foxd3* genomic locus, encompassing its upstream enhancers – E1 – E8, in the Cranial NC (Cl.0, scATAC-seq) and NMps (Cl.5, scATAC-seq) clusters. Black arrow marks a peak specifically accessible in the NMp cluster only, that has not been previously identified as a *foxd3* enhancer. E4 and E6 exhibiting opposite accessibility profiles are marked in pink boxes. (**E**) Lateral view of the whole-mount live *Tg(foxd3:enh4-EGFP)*^*ox110*^ transgenic embryo at 6ss. Left – bright field view; right - Enh4 activity reported by expression of green fluorescence protein (EGFP). Scale bars correspond to 100 μm. (**F**) Lateral view of the whole-mount live *Tg(foxd3: enh6-EGFP)*^*ox109*^ transgenic embryo at 6ss. Left – bright field view; right – Enh6 activity reported by expression of EGFP. Scale bars correspond to 100 μm. (**G**) Bubble plot summarizing fold enrichment and *p-values* (corrected for false discovery rate) for the most significant biological process gene ontology terms associated to differentially expressed genes between foxd3:enh4-EGFP (E4+) and EGFP-(Negs) cells from the same embryo clutches; between foxd3:enh4-EGFP (E4+) and foxd3:enh6-EGFP (E6+) cells; between foxd3:enh6-EGFP (E6+) and EGFP-(Negs) cells from the same embryo clutches; and between foxd3:enh6-EGFP (E6+) and foxd3:enh4-EGFP (E4+) cells. (**H**) Predicted enhancer 6 (Enh6) repressors and enhancer 4 (Enh4) activators in the NMps, and Enh6 activators and Enh4 repressors in the NC. Listed bar plots depict log2 fold change of differentially expressed identified TFs in the Enh6+ vs Enh4+ cells, transcription factor (TF) binding sites of which were enriched within Enh4 (in orange) or Enh6 (in green) DNA sequences.

Additionally, we observed notochord-progenitor cluster (*noto*, *shh*, *foxa2*) near the NMp cluster (fig. 1 A, S1 B) and a ‘Mixed NC’ cluster between the NC and NMp-related clusters. ‘Mixed NC’ cells expressed both NC signature genes and NMp-related genes, particularly, pro-neural factors (*alcama/b*, *chrd*, *foxj1a*) (fig. 1 B, S1 B). The mixed NMp-NC signature was also uncovered in three clusters containing naïve, transitioning cells with lower unique molecular identifier (UMI) counts (Cl.0 – dual/ naïve signature, Cl.1 – more NMp-pronounced, Cl.2 – more pro-neural NC-pronounced) (fig S1 D-F), suggesting a common origin for NMp and pro-neural NC.

Single-cell assay for transposase-accessible chromatin using sequencing (scATAC-seq) (Satpathy et al., 2019) of the citrine+/foxd3+ population (fig S2 A-C) followed by filtering and clustering (Stuart et al., 2020), uncovered 10 specific scATAC clusters (fig. 1 C), which we annotated by transferring labels from appropriately anchored scRNA-seq clusters (fig. 1 A). By integrating both datasets into a single reference, we confirmed that chromatin accessibility correlated with associated gene expression (fig. S2 D). Accordingly, we identified cranial NC (Cl.0) and tail NMp-related cells (Cl.5) with distinct genomic locus accessibility patterns (fig. 1 C).

We hypothesized that distinct enhancers drive *foxd3* expression in cranial NC versus NMps. To test this hypothesis, we examined chromatin accessibility of the *foxd3* genomic locus for these two clusters. Of 8 enhancers within the *foxd3* gene locus (fig. S3 and (Lukoseviciute et al., 2018)), Enh4 located ~70kb upstream of *foxd3* and an element ~8kb upstream of Enh4 were accessible within the NMp cluster and NMp-derived Paraxial mesoderm, but not cranial or trunk NC cells (fig. 1 D, S2 E). On the other hand, Enh6, located ~112kb upstream of *foxd3* was accessible in all NC cell clusters, including cranial and trunk NC, but not in the posterior NMp cluster (fig. 1 D, S2 E). Notably, both enhancers were accessible in the ‘Mixed NC’ population, suggesting a gradual NMp chromatin signature loss and NC signature acquisition in this putatively transitioning cell type (fig. S2 E).

Foxd3 ChIP of foxd3+ cells from embryos between 75% epiboly to 14-16 somite stages showed that both *foxd3* enhancers were directly bound by foxd3, suggesting autoregulation (fig. S3A). Intriguingly, Enh6 remained accessible throughout these stages, whereas Enh4 became compacted by 16ss in the foxd3+ NC cells (fig. S3 A-B). On the other hand, Enh4 but not Enh6 was predominantly accessible in foxd3-naïve cells of early gastrula (50% epiboly stage) (fig. S3 C) hinting that Enh4 is active prior to NC induction. Of note, foxd3 overexpression led to increased Enh6 accessibility and a decreased Enh4 accessibility in early gastrula embryos. This overall suggests that a high concentration of foxd3, such as in specified NC cells, leads to Enh4 compaction (fig. S3 C). Together, these data suggest bimodal autoregulation of foxd3 that involves Enh4 in NMps and Enh6 in NCs.

To directly investigate the spatiotemporal activity of Enh4 and Enh6, we established stable transgenic lines *Tg(foxd3:enh6-EGFP)*^*ox109*^ and *Tg(foxd3:enh4-EGFP)*^*ox110*^ using our enhancer-reporter system (Chong-Morrison et al., 2018). While both elements were active *in vivo* (fig. 1 E-F), they exhibited remarkable spatial differences at 5-8ss. Enh4 was active in trunk regions: tailbud and developing somites (fig. 1 E), whereas Enh6-EGFP reporter was expressed in cranial regions (fig. 1 F). Bulk RNA-seq of Enh4-EGFP+ and Enh6-EGFP+ cells at 5-6ss (fig. S4 A) revealed relatively high expression of posterior *hox* genes in Enh4+ cells, confirming their posterior identity (fig. S4 B-C). In addition, Gene Ontology (GO) terms associated with genes enriched in the Enh4-EGFP+ cells supported their NMp identity, as the topmost terms referred to formation of mesoderm derivatives, such as somites and notochord, as well as spinal cord patterning and somatic motor neuron differentiation (fig. 1 G, S4 B-C). In contrast, Enh6-EGFP+ cells, were depleted for posterior *hox* gene expression and enriched for *bona fide* NC markers (*tfap2a*, *pax3a*, *sox10*) (fig. S4 B,D). Accordingly, the GO terms for this population were associated with NC differentiation (i.e., neural crest, iridophore and enteric nervous system (ENS) development), as well as CNS development (forebrain or fourth ventricle) (fig. 1 G, S4 B,D). Importantly, *foxd3* expression was higher in the Enh6-EGFP+ cells versus Enh4-EGFP+ cells, consistent with our scRNA-seq showing that *foxd3* was expressed at lower levels in NMps and hinting at the functional significance of the differential enhancer usage (fig. S4 B, 1 B). Notably, later in development, Enh6 is active across the entire A-P axis (fig. S4 E). This overall indicates that Enh6 is a *bona fide* NC enhancer.

ATAC-seq of FAC-sorted Enh6-EGFP+ cells (cranial) and Enh4-EGFP+ cells (caudal) at 5-6ss showed that Enh4 and the upstream caudal-specific element (fig. S5 A, black arrow) were only accessible in the cognate, posterior (Enh4+) cells, and not in cranial (Enh6+) cells. Conversely, Enh6 and other NC *foxd3* enhancers, such as Enh1 and Enh5, were only open in the Enh6+ cells, but not in the Enh4+ cells (fig. S5 A). Overall, our data suggest that a mutually exclusive set of *cis*-regulatory elements drives *foxd3* transcription in tail NMp cells compared to cranial NC cells to ensure that the correct level of foxd3 is produced in the distinct cell types.

We integrated TF binding site (TFBS) analysis of each enhancer with RNA-seq of Enh4-EGFP+ and Enh6-EGFP+ cells (fig. 1 H) to uncover potential enhancer-specific upstream regulators (fig. S5 B-C). Our data suggested that NC factors, such as, tfap2a and sox10, activate Enh6 in Enh6-EGFP+ NC cells, whereas posterior *hox* genes repress Enh6 in Enh4-EGFP+ NMps (fig. 1 H, S5 B-C). 5’ *Hox* genes are known to exhibit posterior prevalence via more anterior gene repression (Lewis, 1978; Zhu et al., 2017). Our data further suggested that Enh4 may be activated by NMp regulators such as sall4 and raraa (Gouti et al., 2017; Tahara et al., 2019) in NMp cells and repressed by high levels of foxd3 and other NC regulators, such as zic2b and klf7a, in NC cells (fig. S3, S5 B-C).

We infer that low levels of foxd3, regulated by Enh4, enable the development of NMp prior to their differentiation into trunk NC, whereas high levels of foxd3, regulated by Enh6, promote the NC fate. Mutually exclusive foxd3 autoregulation in NC and NMps and the identification of a foxd3-expressing single-cell cluster with an NMp-NC mixed epigenetic and transcriptional signature (‘Mixed NC’), further imply a relationship between NMp and trunk NC programmes *in vivo*.

### Developmental trajectories of zebrafish NMps

NMp differentiation has been studied in amniotes and *in vitro*, but little is known about NMp development in anamniotes. Although a tailbud *brachyrury*+(*tbxta*+)/*sox2*+ NMp pool capable of giving rise to both mesodermal and neural fates was described in zebrafish (Attardi et al., 2018; Martin and Kimelman, 2012), it remains unclear whether tailbud NMps in zebrafish are bipotent or mono-fated, self-renewing or quiescent, and to what extent they contribute to posterior axis construction. Unlike in amniotes, it has been proposed that the zebrafish NMp pool quickly segregates into mono-fated mesodermal or neural cells, with only a few quiescent NMp-like cells remaining in the tailbud (Attardi et al., 2018).

To study the population of foxd3+ NMp cells we have discovered and its link to NCs, we focused on specific scRNA-seq clusters from our initial analysis, including all NMp-related clusters, the non-cranial/trunk NC (*otx1*-, *otx2b*-) and the transitioning ‘Mixed NC’ cells (fig. 1 A). The increased resolution yielded 7 distinctive sub-clusters: NC (*sox10*, *tfap2a*), ‘Mixed NC’ (*tfap2a*, *cdx4*, *msgn1*), Neural (*alcamb*, *plxna3*), NMp-neural (*sox2*, *sox3*), NMps (*tbxta*, *cyp26a1*), Mesodermal progenitor cells (MPCs) (*tbx16/l*, *msgn1*, *myf5*) and Paraxial mesoderm (*myl10*, *unc45b*) clusters (fig. 2 A, S6 A).

**Figure 2:**
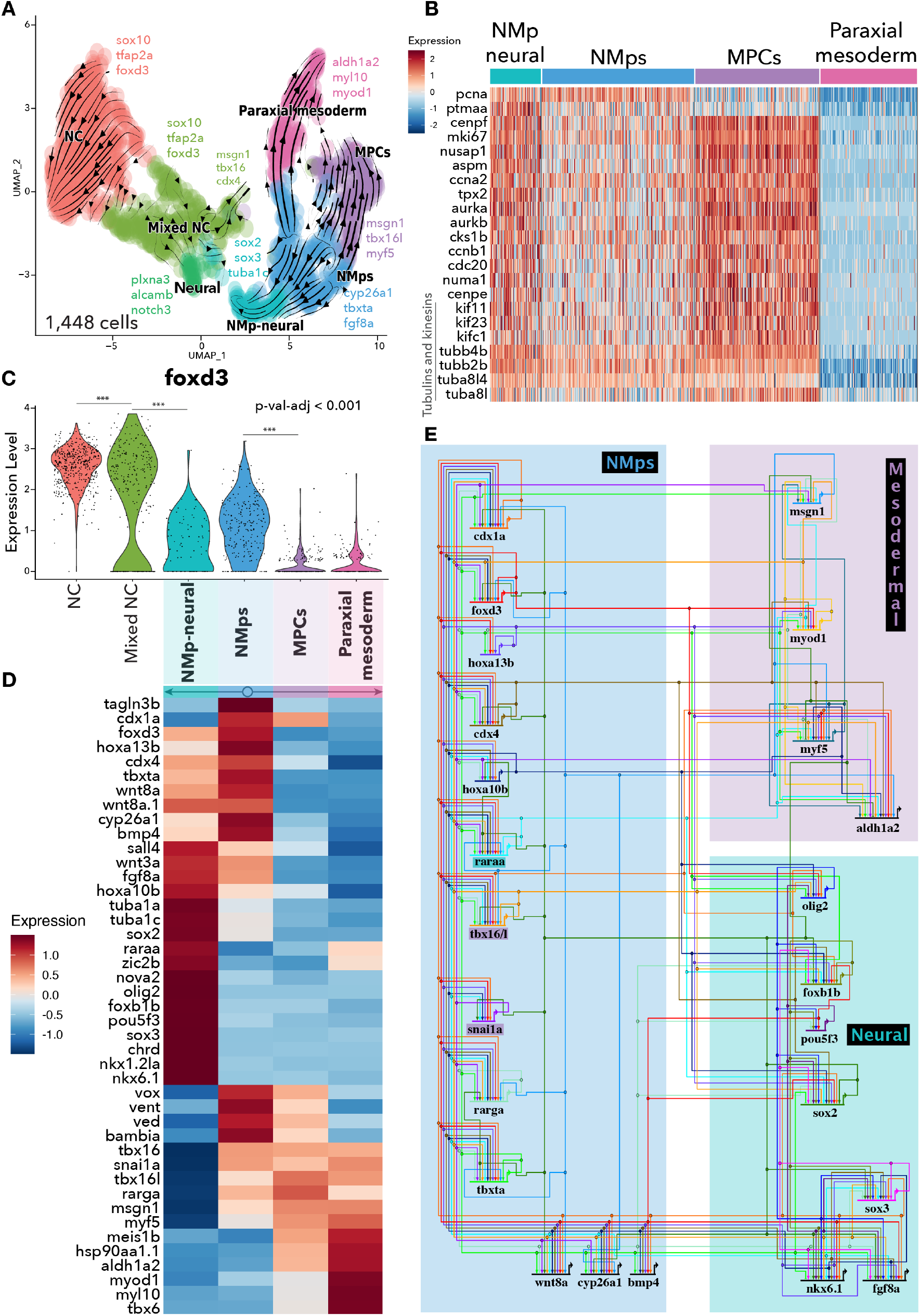
Gene dynamics and gene regulatory networks underlying tailbud neuromesodermal progenitor (NMp) differentiation. (**A**) UMAP embedding showing 7 sub-clusters of foxd3+ 1,448 single cell transcriptomes of selected posterior scRNA-seq clusters: NMps/NMp mesoderm, Paraxial mesoderm, NC-1 and ‘Mixed NC’ from the initial full dataset clustering results (fig. 1 A). Sub-clusters were manually annotated based on the enriched cell-type markers (S8). NC – neural crest, NMps – neuromesodermal progenitors, MPCs – mesodermal progenitor cells. Arrows correspond to averaged cell velocities based on RNA splicing kinetics using the scVelo tool. (**B**) Heatmap displaying cell cycling gene expression across NMp-related cell clusters. (**C**) Violin plots depicting foxd3 expression across different single cell clusters. Each dot corresponds to a single cell expression value. Stars indicate statistically significant difference of *foxd3* expression between clusters (Wilcoxon rank sum test between each cluster). (**D**) Heatmap illustrating selected cluster from (**A**) enriched gene average expression. (**E**) Principal gene regulatory networks in NMps, MPCs and NMp-neural cells reconstructed by picking each scRNA-seq cluster enriched genes as an output and enriched transcription factor binding sites, within associated gene *cis*-regulatory elements from the scATAC-seq, as an input.

We employed the RNA velocity tool, scVelo (Bergen et al., 2020) to determine whether the zebrafish NMp population that we identified at 5-6ss is bipotent. This approach allowed us to dynamically model single cell transcript splicing, transcription and degradation events and predict cellular transitions and latent time. We found that, like their amniote counterpart, zebrafish tailbud NMps are bipotent and give rise to both neural and mesodermal fates at 5-6ss (fig. 2 A).

Furthermore, latent time analysis suggested that, at 5-6ss, NMp cells were the ‘oldest’ cells in the dataset, whereas NMp-derived mono-fated cells, namely NMp-neural and MPC cells, were ‘younger’, more recent populations (fig. S6 B). Therefore, although we detected mono-fated NMp-related cells, the RNA velocity trajectories and predicted cell age suggest that these cells were not allocated early in gastrulation but rather descended from the posterior bipotent NMps, hence contradicting previous claims issued from the analyses performed at later stages of development (21ss) (Attardi et al., 2018). Additionally, bipotent and mono-fated NMp-neural cells and MPCs expressed cell cycling genes at relatively high levels in comparison to differentiated Paraxial mesoderm cells (fig. 2 B), suggesting that they are not quiescent, at least at 5-6ss. *Hox* gene expression in these cell populations further support that NMps reside at the most posterior end of an embryo (*hoxa13b*), whereas the emerging NMp-neural and MPC cells lie slightly more anterior to the NMps (*hoxa11b*) together with even more anteriorly placed Paraxial mesoderm (*hoxb10a*) (fig. S6 C).

### Characterizing gene expression dynamics driving NMp transitions

Next, we analyzed the scRNA-seq data to infer the underlying gene dynamics during NMp transitions in zebrafish model. As seen above, *foxd3* was expressed at lower levels in the NMps than in NC, and its expression was further decreased upon NMp differentiation into MPCs and Paraxial mesoderm (fig. 2 C). It is noteworthy that the ‘Mixed NC’ cluster, which contains transitional NC/NM progenitors, exhibited a bimodal distribution of *foxd3* expression (fig. 2 C). Similar to other studies of NMps, we were unable to pinpoint a single NMp-specific marker (fig. 2 D, S6 D). Genes expressed in NMps (e.g., *cdx1a/4*, *tbxta*, *msgn1*, 5’ *Hoxs*) revealed evolutionary conservation of the NMp Gene Regulatory Network (NMp GRN) across taxa (fig. 2 D, S6 D) (Gouti et al., 2017). Moreover, anamniote bipotent NMps also highly expressed *bmp4* and its inhibitor *bambia* (Paulsen et al., 2011), *tagln3b*, and the dorsoventral (DV) axis patterning genes *vent*, *vox* and *ved* (fig. 2D, S6 D). *Sox2* was expressed at low levels in NMps and was upregulated in NMp-neural cells. Zebrafish NMps highly expressed *cyp26a1*, which is known to guard NMp bipotency via retinoic acid (RA) degradation in the amniotic tailbud (Gouti et al., 2017; Martin and Kimelman, 2012).

Amniote NMps transit to presomitic mesoderm via intermediate mesodermal progenitor cells by first further upregulating *Msgn1* and *Tbx6* genes and then downregulating *Bra* and *Nkx1.2* genes (Gouti et al., 2017). Our data suggest that in zebrafish the role of *tbx6* is replaced by *tbx16/l* homologs, as *tbx6* was highly upregulated only in paraxial mesoderm cells (fig. 2 D, S6 D). Upon mesoderm induction, the genes encoding the RA synthesising enzyme, *aldh1a2*, and retinoic acid receptor (*rarga*) are also upregulated. In amniotes, mesoderm-produced RA feeds back onto the tailbud NMp pool, where *Bra* becomes inhibited and *Sox2* expression becomes upregulated, tipping the NMp balance towards neural differentiation (Gouti et al., 2017). Our data suggest that RA-signalling is controlled in a very similar manner as in amniotes and thus was also conserved, even though in zebrafish RA inhibition is not essential for trunk development as opposed to amniote body axis extension (Berenguer et al., 2018). Upon NMp differentiation towards neural fate, pro-mesodermal genes were downregulated, while genes, such as, *tuba1a/c*, *sox11a*, *sox2*, *pou5f3*, *apela* and *fgf8a*, that were also expressed by NMps, were upregulated (fig. 2 D, S6 D). These cells also switched other neural markers on, such as *sox3*, *notch3*, *alcamb*, *foxj1a/b*, *neurog1*, *nkx1.2la*, *elavl3* and *nova2* upon their differentiation from the NMps to the neural fate (fig. 2 D, S6 D). A prevailing feature of neural induction is Bmp inhibition (Verrier et al., 2018). Correspondingly, we observed a decrease of *bmp4* expression and simultaneous upregulation of the bmp-inhibitor, *chordin*, in NMp-neural cells (fig. 2 D, S6 D).

### Gene regulatory networks (GRNs) underlying anamniote tailbud NMps

To reconstruct the principal GRNs governing NMp development in zebrafish, we identified upstream inputs and downstream effectors by integrating NMp and NMp-derivative gene expression lists and associated *cis*-regulatory elements from the scATAC-seq NMp-specific cluster (fig. S7 A). We extracted all the scATAC peaks specifically found in the NMp cells from the Loupe cell browser (cl.2 from the *k*-mean clustering, fig. S7 B-F), annotated them to the nearest genes, and focused on genes known to be enriched in either NMps, NMp-neural cells or MPCs (fig. 2 D, S6 D). Next, we identified and annotated significantly enriched TF motifs (P<0.0001) from the Hocomoco v11 database, which contains extensive human and mouse TF binding models (Kulakovskiy et al., 2018), using Homer across these three groups and visualised built circuits of the selected core factors using the BioTapestry tool (Longabaugh et al., 2009) (fig. 2 E).

The reconstructed GRN hierarchies in NMp revealed that posterior homeobox genes, *cdx1a*, *cdx4*, *hoxa13b* and *hoxa10b*, as well as *foxd3*, were positioned at the top of the GRN (fig. 2E). *Tbxta* was regulated with inputs from cdx, foxd3, posterior hox, retinoic acid receptor and sox2 factors (fig. 2E). Importantly, regulatory elements controlling *sox2* activity were significantly enriched for tbxta inputs suggesting that, like in amniotes, sox2 and tbxta exerted cross-repressive transcriptional activity (Gouti et al., 2017). Such cross-repression was also apparent between sox2 and tbx16/l regulation. *Msgn1* emerged on the top of the mesodermal compartment hierarchy with the upstream inputs, such as tbxta, tbx16/l, snai1a and retinoic acid receptors (fig. 2 E). Conversely, the neural fate drivers were olig2, foxb1b and pou5f3, regulating each other and the downstream neural targets (fig. 2 E), whereas their upstream regulators appeared to be posterior homeobox, retinoic acid receptor and foxd3 factors. *Foxb1b* and *olig2 cis*-regulatory elements were also marked by tbxta and tbx16/l motifs, suggesting transcriptional inhibition of the neural fate within the pro-mesodermal circuit (fig. 2 E). Overall, by integrating our scRNA-seq and scATAC-seq, we reconstructed a comprehensive multi-circuit GRN governing NMp differentiation, precisely detailing direct interactions and hierarchical organisation within the network. This GRN is poised to inform future direct differentiation-based neural cell therapies.

To validate these findings we used the hybridisation chain reaction (HCR) to visualise transcripts identified as enriched in NMp-related clusters (fig. 2 D, S6 D, 3 A-B) in whole mount embryos (Choi et al., 2018). Our HCR results localised *tbxta*+/*sox2*+ NMps to within the dorsal part of the tailbud, matching previous studies in zebrafish (Attardi et al., 2018; Martin and Kimelman, 2012). *Enh4-EGFP* transcripts overlapped the posterior *tbxta*+/*sox2*+ NMp territory but were not restricted to it (fig. 3 C-C”). Interestingly, the more anterior dorsal tailbud region was positive for *sox2* and *Enh4-EGFP* but not *tbxta* transcripts, whereas the ventral part of the tailbud was positive for *tbxta* and *Enh4-EGFP* but not *sox2* transcripts (fig. 3 C’-C”). Therefore, our results suggest that *Enh4*+ cells comprise not only a bipotent *tbxta*+/*sox2*+ NMp population, but also NMp-derived early neural and mesodermal progenitors.

**Figure 3:**
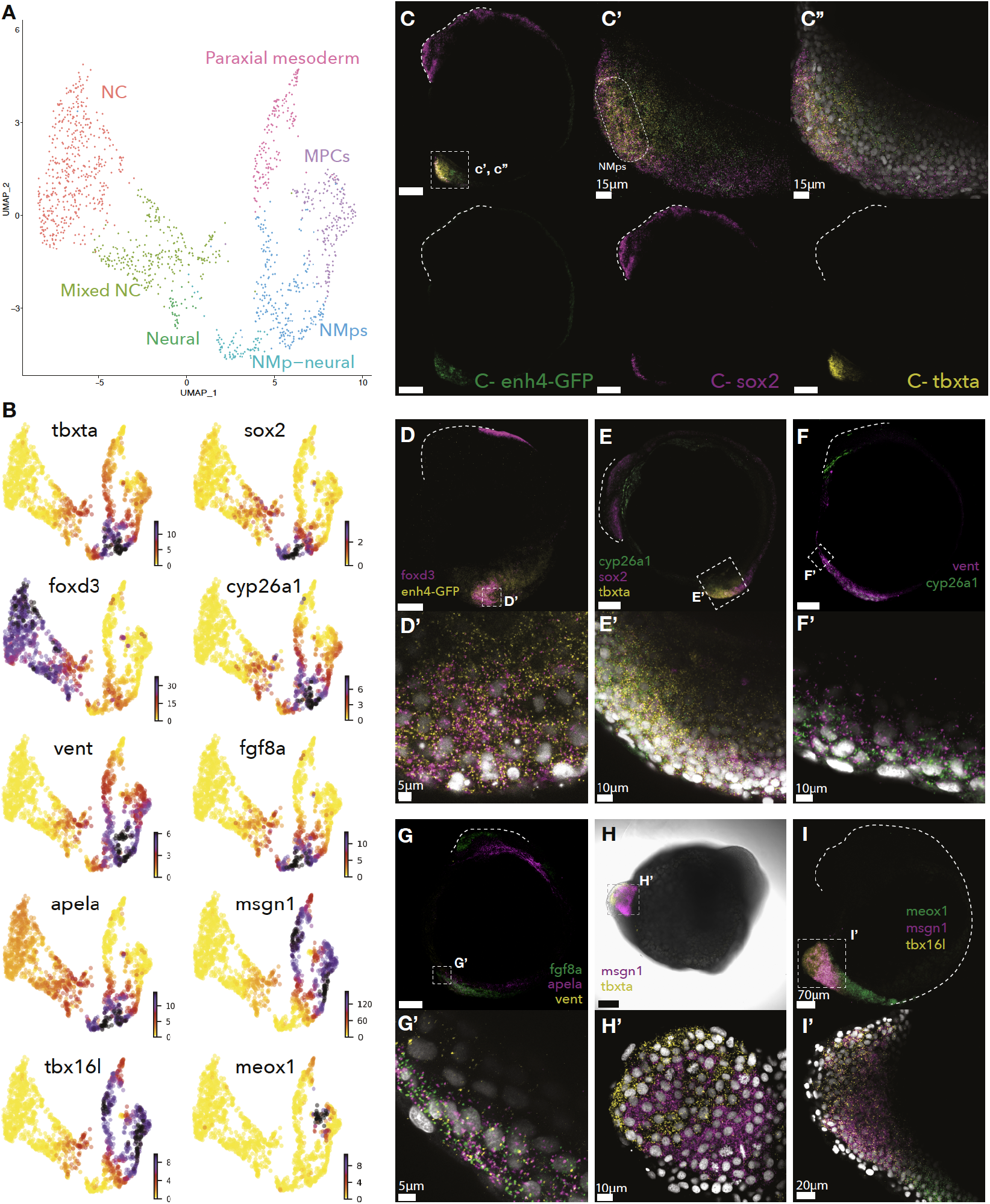
Spatial validation of zebrafish tailbud neuromesodermal progenitor (NMp) and early derivative expressed genes. All embryos are 5-6 somite stage (ss) and in lateral view, unless stated otherwise. Not specified scale bars correspond to 100 μm. (**A**) UMAP embedding showing 7 sub-clusters of foxd3+ 1,448 single cell transcriptomes. NC – neural crest, NMps – neuromesodermal progenitors, MPCs – mesodermal progenitor cells. (**B**) (**C**) *In situ* hybridisation chain reaction (HCR) of *Tg(foxd3:enh4-EGFP)*^*ox110*^ transgenic embryo showing EGFP and endogenous *sox2* and/or *tbxta* transcript localisation. (**C’-C”**) Zoomed-in tailbud region from (**C**) showing EGFP transcript and endogenous *sox2* and/or *tbxta* transcript co-localisation and close proximity to single nuclei (**C”**, in grey). (**D**) HCR of *Tg(foxd3:enh4-EGFP)ox110* transgenic embryo and (**D’**) zoomed-in tailbud region showing *EGFP* and *foxd3* transcript co-localisation and close proximity to single nuclei (in grey). (**E**) HCR of wild-type (WT) embryo and (**E’**) zoomed-in tailbud region showing *cyp26a1*, *sox2* and *tbxta* transcript co-/localisation and close proximity to single nuclei (in grey). (**F**) HCR of WT embryo and (**F’**) zoomed-in tailbud region showing *cyp26a1* and *vent* transcript co-/localisation and close proximity to single nuclei (in grey). (**G**) HCR of WT embryo and (**G’**) zoomed-in tailbud region showing *cyp26a1*, *sox2* and *tbxta* transcript co-/localisation and close proximity to single nuclei (in grey). (**H**) HCR of WT embryo (ventral view) and (**H’**) zoomed-in tailbud region showing *msgn1* and *tbxta* transcript co-/localisation and close proximity to single nuclei (in grey). (**I**) HCR of WT 8-10ss embryo and (**I’**) zoomed-in tailbud region showing meox1, msgn1 and tbx16l transcript co-/localisation and close proximity to single nuclei (in grey).

We also observed a population of tailbud cells that co-express *foxd3* transcripts and *Enh4-EGFP* (fig. 3 D-D’), further supporting that Enh4 regulates foxd3 during NMp development. Furthermore, transcripts of RA-degrading enzyme, *cyp26a1*, which was enriched in the NMp cluster (fig. 3 A-B), were observed in the cells that co-express *tbxta* and *sox2* (fig. 3 E-E’), as well as in cells that express the dorsal fate repressor, *vent* (fig. 3 F-F’), which was enriched in the NMp and MPC clusters (fig. 3 A-B). *Fgf8a* and *apela* transcripts, which were enriched in the NMp-neural and NMp clusters (fig. 3 A-B), were also co-expressed with *vent* in the tailbud (fig. 3 G-G’). Similarly, *msgn1* was co-expressed with the NMp gene *tbxta* (fig. 3 H-H’), as well as *tbx16l* (fig. 3 I-I’), identifying cells undergoing an NMp-to-MPC transition (fig. 2 D, S6 D, 3 A-B). Mesodermal homeodomain protein, *meox1*, was expressed anterior to the *msgn1*+/*tbx16l*+ MPCs, including in forming somites (fig. 3 I), with a gradual shift toward anterior from *msgn1*+/*tbx16l*+ /*meox1*-cells via *msgn1*+/*tbx16l*+/*meox1*+ cells and finally *msgn1*-/*tbx16l*-/*meox1*+ differentiated paraxial mesoderm cells (fig. 3 I-I’).

Collectively, our scRNA-seq and HCR data confirm the presence of the NMp pool and its early derivatives at the trunk embryo regions, corresponding to the Enh4+ cells. Furthermore, we have demonstrated clear parallels between the cellular trajectories and transitions (UMAP of clustered single-cells) and localisation of NMps and their derivatives in the embryo (compare fig. 3B and 3C-I). Our results suggest that the tailbud NMps are bipotent and can give rise to neural and mesodermal fates. Accordingly, we have identified gene dynamics and GRNs underlying these transitions.

### Loss of foxd3 leads to the expansion of NMps/derivatives at the expense of transitory pro-neural trunk NC cells

Given the involvement of foxd3 in the development of both NMp and NC, and its cross- and autoregulatory activity along A-P axis possibly linking the two populations, we investigated potential roles for foxd3 in different NMp compartments. To this end, we used an incross between two foxd3 loss-of-function mutant genetrap zebrafish transgenic lines, *Gt(foxd3-citrine)*^*ct110a*^ and *Gt(foxd3-mCherry)*^*ct110aR*^, which each express a truncated and non-functional foxd3 protein fused to either Citrine or mCherry fluorophore, respectively (Hochgreb-Hägele and Bronner, 2013; Lukoseviciute et al., 2018). Using this *foxd3* knock-out strategy, we performed foxd3-(fig. S8 A) alongside the foxd3+ scRNA-seq for comparative analyses.

By integrating/harmonising scRNA-seq datasets from Citrine+/mCherry+ foxd3-cells and from Citrine+/foxd3+ cells into a single reference using Seurat pipeline (Stuart et al., 2019), we defined 17 distinct clusters, including the NC, hindbrain, NMp-related and ‘Mixed NC’ cluster containing transitional NC/NM progenitors (fig. 4 A, S8 B-C). Comparative analysis of foxd3+ (Citrine+) vs foxd3-(Citrine+/mCherry+) cells revealed that loss of foxd3 led to a significant reduction in cells with a canonical NC signature (fig. 4 B), in line with our previous study that reported a decrease in expression of neural crest specification genes in the *foxd3* mutant (Lukoseviciute et al., 2018).

**Figure 4:**
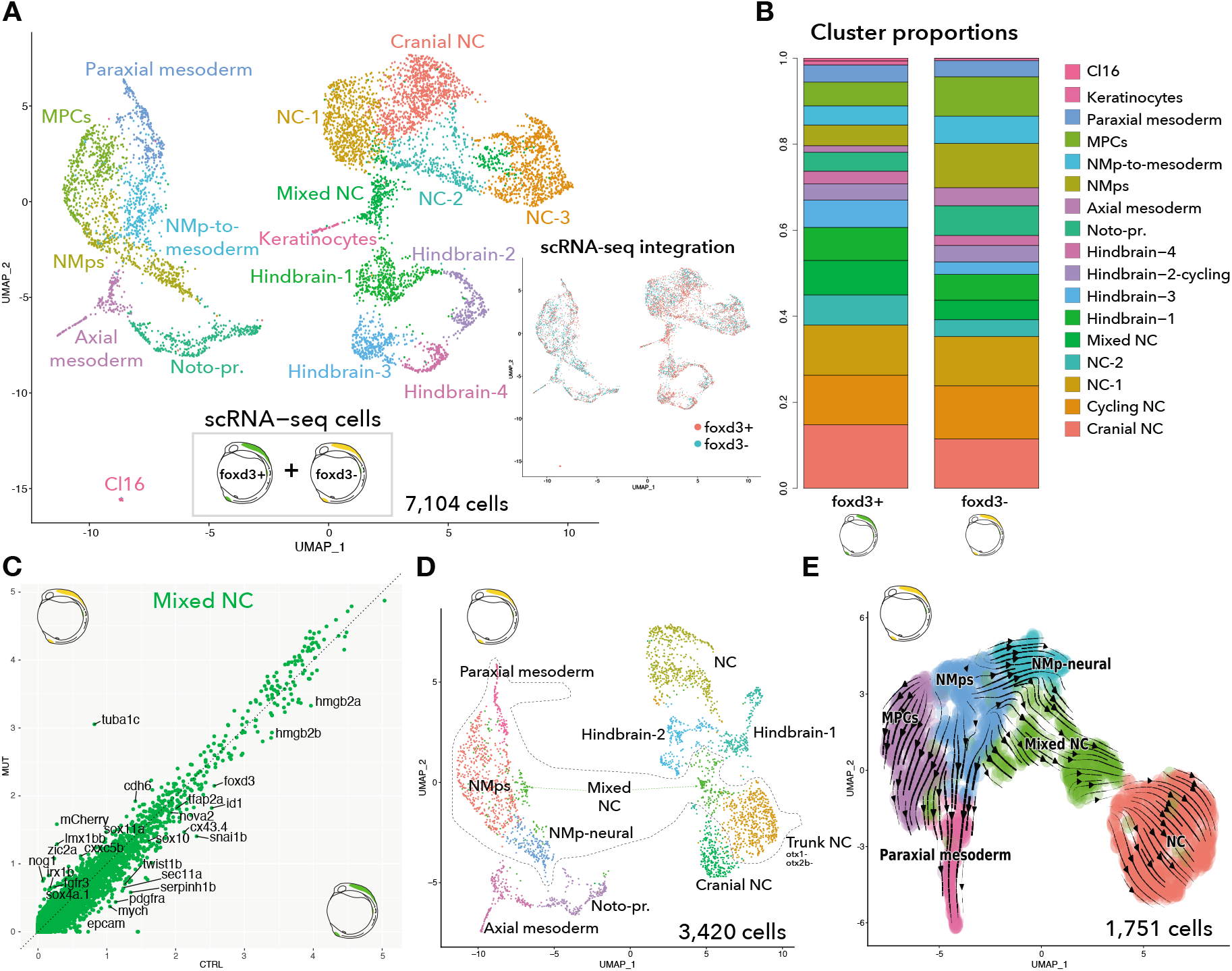
Absence of foxd3 leads to neuromesodermal progenitor (NMp) cell expansion at the expense of neural crest (NC) loss. (**A**) UMAP embedding showing 17 integrated clusters of foxd3+/(citrine+) 3,684 cells and foxd3-(citrine+/mCherry+) 3,420 cells enriched during fluorescence-activated cell sorting (FACS) from 5-6ss embryos of *Gt(foxd3-citrine)*^*ct110a*^ and *Gt(foxd3-mCherry)*^*ct110aR*^ transgenic line in-crosses. Clusters were manually annotated based on the enriched cell-type markers (S10 B). Noto-pr. – notochord progenitors, MPCs – mesodermal cell progenitors, NMps – neuromesodermal progenitors, NC – neural crest. (**B**) Comparative cell number proportions within integrated clusters split between foxd3+ and foxd3-samples. (**C**) Scatter plot displaying average expression of ‘Mixed NC’ cluster genes by both foxd3+/(citrine+, x-axis) and foxd3-(citrine+/mCherry+, y-axis) cells. (**D**) UMAP embedding showing 11 clusters foxd3-(citrine+/mCherry+) 3,420 cells enriched during fluorescence-activated cell sorting (FACS) from 5-6ss embryos of *Gt(foxd3-citrine)*^*ct110a*^ and *Gt(foxd3-mCherry)*^*ct110aR*^ transgenic line in-crosses. (**E**) UMAP embedding showing 6 sub-clusters of foxd3-1,751 single cell transcriptomes of selected initial posterior foxd3-scRNA-seq clusters (marked by dashed outline in D). Sub-clusters were manually annotated based on the enriched cell-type markers (S11 D). Arrows correspond to averaged cell velocities based on RNA splicing kinetics using the scVelo tool.

Notably, the pro-neural ‘Mixed NC’ cluster was also significantly reduced in the *foxd3* mutant (fig. 4 B). NC regulators, such as TFs *tfap2a*, *sox10*, *foxd3*, *twist1b*, cell junction molecules *epcam* and *cx43.4*, and high-mobility group protein genes, *hmgb2a/b*, were expressed at much lower levels in *foxd3*-mutant versus control ‘Mixed NC’ cells (fig. 4 C). Conversely, NMp-neural *tuba1c* and *sox11a* genes (fig. S6 D) and the pro-epithelial-mesenchymal transition (EMT) gene, *cdh6* (Clay and Halloran, 2014) (fig. 4C) were expressed at higher levels in *foxd3*-mutant versus control ‘Mixed NC’ cells. *Tuba1c* and *cdh6* were also upregulated in the mutant NMp and NC-1 clusters (fig. S8 D-E). Strikingly, loss of foxd3 led to an increase in NMp-related cells as well as notochord precursors (fig. 4 B).

Differential analysis between *foxd3*-mutant and *foxd3*-control cells within the NMp-related clusters (fig. S9 A) did not uncover drastic differences in gene expression, with only a few NMp (*tbxta*, *fgf8a*, *cyp26a1*) and mesodermal (*msgn1*, *myf5*, *myod1*) genes downregulated in the mutant cells (fig. S9 A-B). However, pro-neural tubulins (*tuba1a/c*) were up-regulated in the absence of foxd3 (fig. S9 A,C). Therefore, differently from its role in the NC, foxd3 is not critical for NMp induction and development, but it may promote the mesodermal and repress the neural NMp fate.

By scrutinising *foxd3*-mutant cells alone, we uncovered further compositional and dynamical changes within single-cell cluster organisation. To begin with, the number of NC clusters in the mutant was reduced (fig. 4 D) as compared to the control cells (fig. 1 A). Conversely, the NMp-neural cluster in the mutant was more pronounced while ‘Mixed NC’ cluster was distinctly split between the NC and NMp related clusters in the absence of foxd3 (fig. 4 D). Similar to our trajectory analysis of Citrine+/foxd3+ scRNA-seq, we subclustered and reanalysed *foxd3*-mutant cells from NMp-related, ‘Mixed NC’ and non-cranial/trunk NC cluster. The high-resolution cluster structure and gene expression composition (fig. 4 D-E, S9 D-E) was very similar to the equivalent analysis of control cells (fig. 2 A). However, trajectory analysis revealed NMp differentiation into not only MPC and NMp-neural fates, but also towards the ‘Mixed NC’ cluster, which, strikingly, then followed the trunk NC differentiation trajectory (fig. 4 E). Although global velocity analysis in the control condition at 5-6 ss failed to identify NMp-NC trajectories, we revisited the trajectory analysis using a selected cell route tracing approach and were able to demonstrate a cell fate link of NMp-neural and non-cranial/trunk NC cell populations (fig. S9 F).

In the absence of foxd3, NC specification at 5-6ss was perturbed, thus resulting in a reduced number of NC cells, including the pro-neural ‘Mixed NC’ cluster that gives rise to trunk NC, whereas the naïve NMps were expanded. Dynamic RNA velocity analysis of *foxd3*-mutant cells revealed a transition of NMps towards the ‘Mixed NC’ and trunk NC populations that was not detected in the control condition. It is plausible that the NMp population was expanded at the expense of the transitioning NMp-NC cells in the *foxd3*-mutant because it was retained in a more progenitor state at 5-6ss and thus delayed in its typical trajectory. Therefore, the analysis of the NMp trajectory in the *foxd3*-mutant has likely unmasked an NMp-NC progenitor transition that takes place earlier in development that is dependent on the presence of foxd3. This highlights a conserved foxd3 role in specifying NC across the A-P axis.

### Posterior NMp-neural cells give rise to pro-neural NC cells

Analyses of the NMp developmental trajectories in *foxd3*-mutant cells hinted at a possible NMp to posterior pro-neural NC transition, likely taking place prior to the 5-6ss embryonic stage. To analyse the NMp/descendent cell transcriptomes at earlier stages, we isolated Enh4-EGFP+ cells from *Tg(foxd3:enh4-EGFP)*^*ox110*^ embryos at 90% epiboly and bud stages (end of gastrulation/start of neurulation) and analysed them by scRNA-seq. After merging both samples into a single reference (fig. S10 A-B), we uncovered 9 clusters: two NMp clusters, NMp-neural, MPCs, three paraxial/somatic mesoderm clusters, notochord precursors and, importantly, an NC-like cluster (fig. 5 A-B, S10 C-D). NMp-1 and NMp-2 cluster cells expressed similar markers, with the NMp-2 showing a more pronounced NMp signature resembling 5-6ss tailbud NMps (fig. 2 D, S11 A), while NMp-1 in addition expressed chromatin organisation components (*h3f3b.1*, various high mobility group genes) at high levels (fig. 5 B, S10 D).

**Figure 5:**
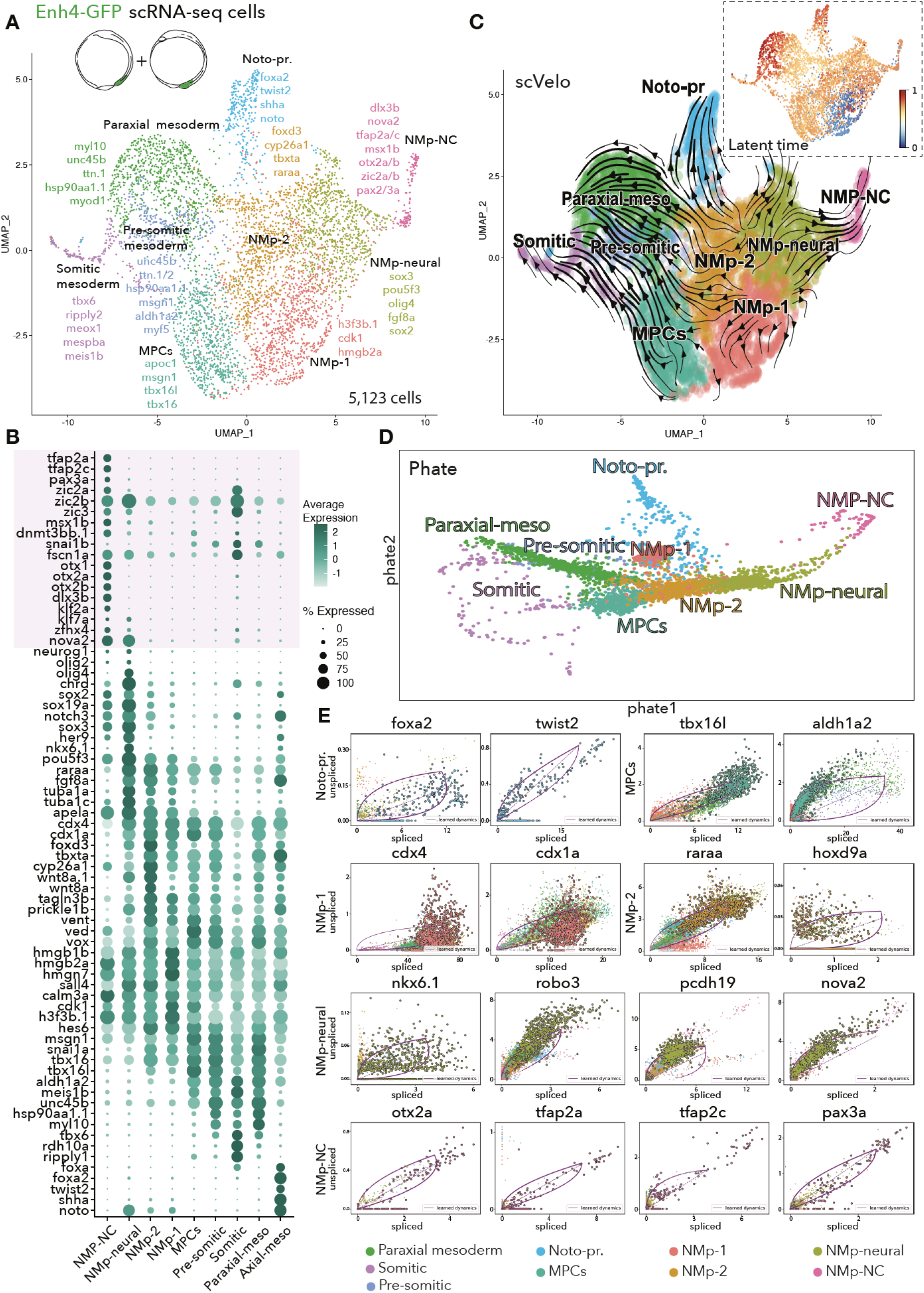
Pro-neural tailbud neuromesodermal progenitors (NMps) give rise to pro-neural trunk neural crest (NC) at bud stage. (**A**) UMAP embedding projecting integrated 9 clusters of 5,123 cells enriched during fluorescence-activated cell sorting (FACS) from *Tg(foxd3:enh4-EGFP)*^*ox110*^ transgenic embryos at 90% epiboly and 1 somite-stage. Clusters were manually annotated based on the enriched cell-type markers (S12 D). NMp – neuromesodermal progenitor, NC – neural crest, MPCs – mesodermal progenitor cells, Noto-pr. – notochord progenitors. (**B**) Dotplot illustrating selected cluster from (**A**) average expression and percentage of cells expressing NC, NMp and NMp-derivative associated genes. Pink box marks neural plate border and early NC genes enriched in the NMp-NC cluster. (**C**) Cluster trajectory analysis; arrows correspond to averaged cell velocities based on RNA splicing kinetics using the scVelo tool. Relative cell age inferred by scVelo latent time analysis is displayed in the top right box. (**D**) PHATE embedding to visualise trajectory branches of (**A**) clusters. (**E**) Predicted driver genes for Noto-pr., MPC, NMp-1/2, NMp-neural and NMp-NC transitions inferred by scVelo dynamical modelling high likelihoods.

We also identified a cluster expressing notochord genes (*noto*, *foxa2*, *shha*) (fig. 5 B, S10 D). Therefore, it is likely that the Enh4-EGFP+ posterior population includes not only NMps, but also earlier blastoderm margin cells that give rise to notochord. Moreover, a recent study in chicken has revealed that the anterior part of the primitive streak contributes to NMp development in addition to the notochord (Guillot et al., 2020; Tam and Beddington, 1987).

We uncovered multiple pro-mesodermal clusters with a progressive loss of MPC markers (*tbx16/l*, *tbxta*, *msgn1*) and acquisition of mature mesoderm markers (*myl10*, *unc45b*) (fig. 5 B, S10 D). As at 5-6ss, we identified an NMp-neural cluster expressing NMp (*msgn1* and *tbx16l*) and pro-neural genes (*sox2/3*, *olig4*, *tuba1a/c*, *alcamb*) (fig. 5 B, S10 D). However, at this earlier stage, we also identified a cluster sharing a number of NMp-neural cluster markers (*cdx4*, *msgn1*, *nova2*, *sox2*), but also expressing neural plate border/early NC markers (*pax3a*, *tfap2a/c*, *msx1b*, *zic2a/3*) (fig. 5 B, S10 D). Curiously, this NC-like cluster did not express high levels of *foxd3* (fig. 5 B). We predicted Enh4 was repressed by TFs, such klf and zic (fig. 1 H), which are upregulated upon NC specification. Indeed, our analysis of transcriptional dynamics during NMp-NC specification shows that these repressive factors were increased (fig. 5 B), which in turn likely leads to *foxd3* enhancer (Enh4) decommissioning and hence loss of *foxd3* expression. Shortly after expression of NC factors, such as *tfap2a* and *sox10*, *foxd3* Enh6 is activated that subsequently upregulates *foxd3* expression in the developing NC cells (fig. 1 H).

Similar to trajectory analysis at 5-6ss, NMp-1/2 velocities indicated transition towards mesodermal and NMp-neural clusters (fig. 5 C). However, we were now also able to detect prominent velocities from NMp-neural to NMp-NC cluster (fig. 5 C). Of note, NMp-NC, NMp-neural, MPCs, NMp-1/2 and somitic cell clusters expressed cell cycling genes (fig. S11 B), suggesting they were highly proliferative at the bud stage. Furthermore, whereas NMp-2 and NMp-neural cells expressed posterior *hox* genes, NMp-NC cells did not, possibly due to a lost spatial identity during migration (fig. S11 C). To solidify our findings, we compared our scVelo results to results obtained using other available cell lineage/pseudotime analysis tools, such as Slinghshot and Phate. These approaches are based on lineage identification using minimum spanning tree as well as pseudotime and data diffusion geometry for dimensionality-reduction, respectively (Moon et al., 2019; Street et al., 2018). Both tools agreed with scVelo showing NMp-neural cells giving rise to the NMp-NC cluster (fig. 5 C-D, S11 D). Phate embedding placed NMp-1 cluster adjacently to both Notochord precursors and NMp-2 cluster (fig. 5 D). Additionally, based on latent time analysis, the NMp-1 cluster was predicted to be the ‘oldest’ cluster of the Enh4-EGFP+ cell population (fig. 5 C), suggesting that the NMp-1 cluster corresponds to the remnant cells of the blastoderm margin giving rise to NMps and notochord precursors. Consistently with scVelo, Phate placed NMp-2 cluster between NMp-neural and MPC cells, suggesting NMp-2 cluster is able to give rise to both mesodermal and neural fates and hence it corresponds to the tailbud NMps (fig. 5 D).

To further understand which genes drive NMp developmental transitions, we again employed the dynamical scVelo modelling (see fig. S11 D for each cluster top 10 likelihood genes). As intronic reads are crucial for computing learned dynamics for each gene, we were not able to examine single exon or single short intron gene dynamics of *foxd3* or *sox2*. Our analyses showed that *foxa2* and *twist2* contributed to driving notochord precursor transitions, the NMp-1 cluster transitions were driven by posterior *cdx* genes, while posterior *hox* genes and retinoic acid receptor (*raraa*, *rarga*) expression were steering NMp-2 cluster transitions (fig. 5 E). Previously uncovered pro-mesodermal (*tbx16l*, *aldh1a2*) and pro-neural (*nova2*, *nkx6.1*) genes were contributing to the corresponding transitions, whereas a number of the pro-neural genes (*nova2*, *pcdh19*) contributed to the NMp-NC transitions, which were additionally directed by NC genes, such as *tfap2a/c* or *pax3a* (fig. 5 E).

To validate our Enh4-EGFP+ scRNA-seq results, we employed HCR to visualise multiple transcripts in 1-4ss whole-mount embryos. *Foxd3* expression patterns revealed that in addition to its anterior expression, *foxd3* was co-expressed in the tailbud with the NMp gene, *tbxta*, and Enh4-driven EGFP in the bud stage embryos, once again confirming its involvement in the tailbud NMp development (fig. 6 A, B-B’). To validate expression of early NC genes in the NMp-related tailbud cells, we performed HCR using probes for *tfap2a* and *pax3a* transcripts, which were enriched in our NMp-NC cluster and contributed to its transition (fig. 5 A-B,E, 6 A). We identified some cells emerging from the tailbud, that co-expressed *enh4-EGFP* and *tfap2a*, with a gradual decrease of *EGFP* and increase of *tfap2a* transcripts away from the midline (fig. 6 C-C’). Similarly, *tfap2a* was co-expressed with the NMp genes, *sox2* and *tbxta* (high *sox2*/low *tbxta* levels) (fig. 6 D-D’) in agreement with our scRNA-seq results (fig. 6 A) and NMp-derived NC pro-neural bias. *Pax3a* expression was detected lateral to the tailbud (fig. 6E), where *Enh4-EGFP*, *foxd3* and *pax3a* transcripts were localised to a number of single nuclei (fig. 6E’). Additionally, we confirmed *pax3a* was co-expressed with *sox2* and *msgn1* transcripts in the anterior/lateral part of a tailbud (fig. 6 F-G’).

**Figure 6:**
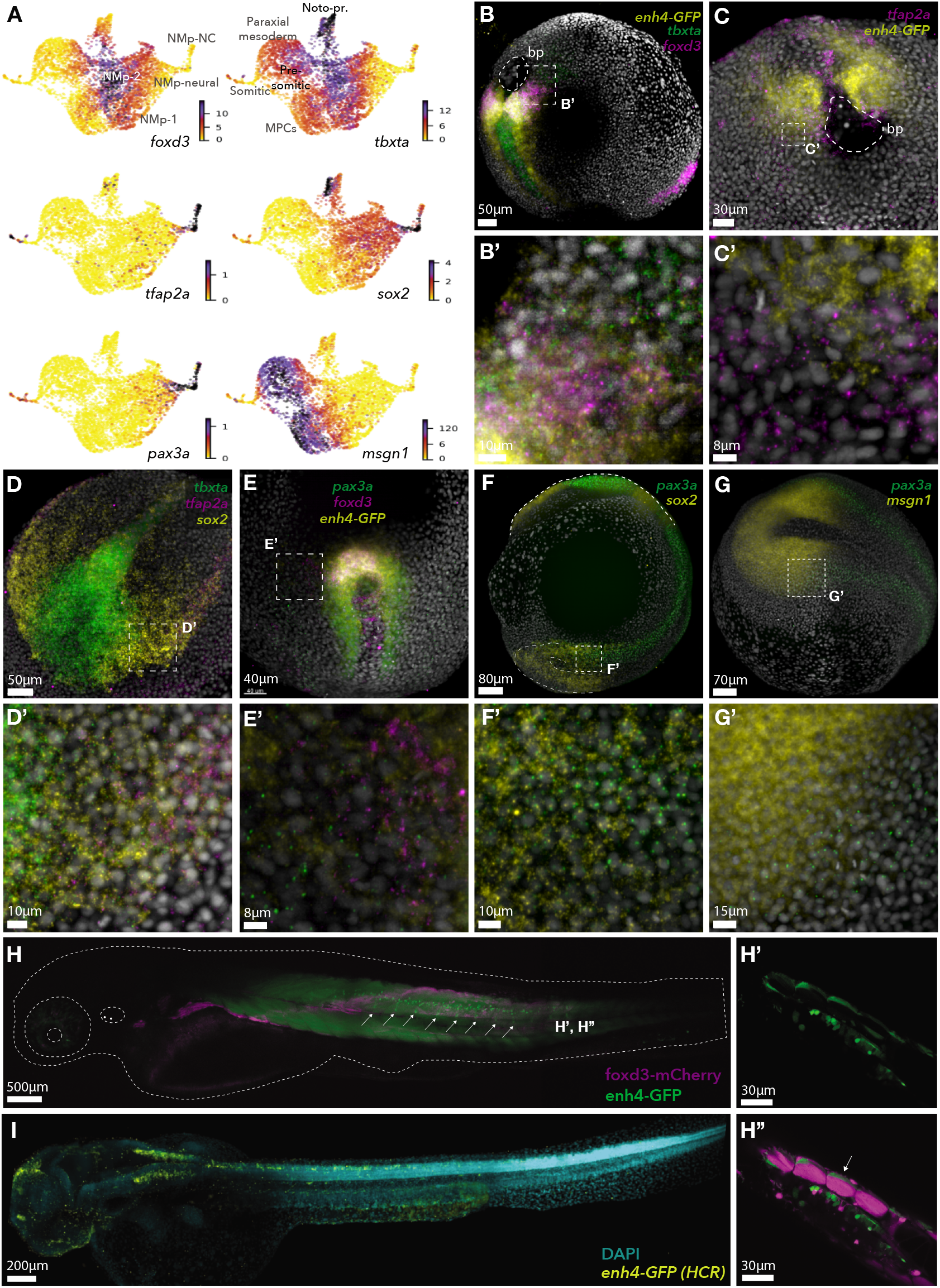
Spatial validation of zebrafish transitory cell population of tailbud neuromesodermal progenitor (NMp)-neural crest expressed genes. All embryos are 1-4 somite stage (ss). (**A**) The expression of spatially validated genes projected on UMAP. (**B**) *In situ* hybridisation chain reaction (HCR) on *Tg(foxd3:enh4-EGFP)*^*ox110*^ transgenic embryo and (**B’**) zoomed-in lateral side of a tailbud showing *EGFP*, *foxd3* and *tbxta* transcript co-/localisation and close proximity to single nuclei (in grey); bp – blastopore. (**C**) Dorsal view of HCR on *Tg(foxd3:enh4-EGFP)*^*ox110*^ transgenic embryo and (**C’**) zoomed-in lateral side of a tailbud showing *EGFP* and *tfap2a* transcript co-/localisation and close proximity to single nuclei (in grey). (**D**) Dorsal view of wild-type (WT) embryo tailbud HCR and (**D’**) zoomed-in lateral tailbud region showing *tbxta*, *sox2* and *tfap2a* transcript co-/localisation and close proximity to single nuclei (in grey). (**E**) Dorsal view of *Tg(foxd3:enh4-EGFP)*^*ox110*^ transgenic embryo tailbud HCR and (**E’**) zoomed-in region lateral to tailbud showing *EGFP*, *foxd3* and *pax3a* transcript co-/localisation and close proximity to single nuclei (in grey). (**F**) HCR of WT embryo (lateral view) and (**F’**) zoomed-in region anterior to tailbud showing *sox2* and *pax3a* transcript co-/localisation and close proximity to single nuclei (in grey). (**G**) Dorsal view of WT embryo HCR and (**G’**) zoomed-in tailbud region showing *msgn1* and *pax3a* transcript co-/localisation and close proximity to single nuclei (in grey). (**H**) Lateral view of 2 days post fertilisation (dpf) zebrafish larva (*Gt(foxd3-mCherry)*^*ct110aR*^ transgenic line crossed with *Tg(foxd3:enh4-EGFP)*^*ox110*^) expressing foxd3-mcherry (coloured in magenta) and Enh4-GFP (in green). Letters H-H’, correspond to an approximate position within a zebrafish of the sections listed on the right. White arrows point to dorsal root ganglia. (**I**) Transcriptional Enh4 activity at 2 dpf inferred by *in situ* hybridisation chain reaction (HCR) using probes against *EGFP* transcript driven by Enh4.

Our results strongly suggest that at least a portion of posterior neural crest derives from the tailbud neuromesodermal progenitor population. The split appears to take place early during development, at the end of gastrulation and before the extension of the embryonic axis. This transition occurs via an intermediary of pro-neural NMps, which fits with a known trunk NC bias for neural lineages. In line with this analysis, in 2-day old zebrafish embryo the Enh4-EGFP signal (NMps) was detected in the posterior neurons and mesoderm, including NC-derived dorsal root ganglia (fig. 6H, H’, H”). The enhancer remained active in these tissues as seen by detection of EGFP transcripts (fig. 6I). Therefore, our result advocate for a shared pro-neural NMp and posterior NC origin *in vivo*.

## Discussion

Cranial and trunk NC exhibit quite different transcriptional and plasticity properties and hence generate distinct derivatives, with the cranial NC yielding both ectomesenchymal and neuronal derivatives, and trunk NC having a clear neuronal bias. Here we provide evidence that divergent embryonic origins underlie this dichotomy. We demonstrate that at least a portion of trunk neural crest cells in the developing zebrafish embryo arise from neuromesodermal progenitors rather than from the neural plate border epiblast. Interestingly, these trunk neural crest progenitors form via an intermediary neural progenitor fate, much like *in vitro* NMps derived from hPSCs that require a passage via pre-neural fate to acquire posterior trunk NC features progressively (Cooper et al., 2020). All human *in vitro* differentiation protocols that successfully generate trunk NC identities must first generate an intermediate neuromesodermal axial progenitor, drive them to a pro-neural fate, in order to form NC (Frith et al., 2018; Gomez et al., 2019; Hackland et al., 2019). Our findings, however, extend beyond the synthetic *in vitro* differentiation systems and unravel underlying gene regulation during simultaneous NMp transitions into mature mesodermal, neural and NC fates within the *in vivo* context of developing embryos. The overall principles we have uncovered in the zebrafish are largely conserved in mouse and hPSC-derived NMp transitions. For instance, the core zebrafish NMp gene battery is fundamentally conserved with the *in vitro* human and mouse NMps (Brachyury, Sox2, Cyp26a1, Cdx4, Wnt8a), although the role of zebrafish foxd3 likely replaced by other Fox family TFs, such as Foxd1, in mouse (Gouti et al., 2017). As a result, our *in vivo* findings may be highly relevant to trunk NC in humans and may represent a general mechanism to be further explored for therapeutic approaches in regenerative medicine.

We discovered that foxd3 is an important regulator of both NC and NMp cell populations. Strikingly, we discovered that foxd3 is driven by different sets of autoregulated enhancers in the posterior NMps and the anterior or trunk NC (fig. 1, S2, S7). Similarly to the *in vitro* human trunk NC differentiation studies (Frith et al., 2018; Gomez et al., 2019; Hackland et al., 2019), we observed a transitioning pro-neural ‘Mixed NC’ cell population exhibiting not only NC, but also NMp gene expression signatures (fig. 1, 2, S1, S10). Notably, in the *foxd3* knock-out conditions, NMp induction was not severely perturbed, whereas NC cell development was largely impaired (fig. 4, S11). Moreover, NMps expanded at the expense of NC cell number deficiency (fig. 4). This indicated an underlying intricate relationship between NMps and NC *in vivo* and highlights the importance of foxd3 during the NMp to NC transition. We further demonstrated that at earlier developmental stages, NMp-neural cells directly transitioned to become future trunk pro-neural NC progenitors (fig. 5, 6). Strikingly, these cells did not express *foxd3* at high levels (fig. 5, 6). Upon NC induction, Enh4 becomes inactivated leading to reduction of *foxd3* transcription. Given that later at 5-6ss we see bimodal (low and high) *foxd3* expression in the ‘Mixed NC’ cluster (fig. 2), we hypothesise that initiation of an NC-specific enhancer, such as Enh6, leads to reactivation of *foxd3* in the developing trunk NC. Therefore, a *foxd3* transcriptional continuum is transiently interrupted during the programme switching.

In this study, we have uncovered that early anamniote tailbud NMps are regulated by evolutionary conserved GRNs capable of giving rise to mesodermal, pro-neural spinal cord and pro-neural trunk NC cells. The extensive datasets and genetic tools presented here may also provide a resource for future experiments aiming to determine to what extent these axial progenitors contribute to the trunk NC formation in zebrafish and across taxa. It will be important to further understand the evolutionary origin and heterogeneity of trunk NC and how we could employ this knowledge to stir axial progenitor transitions into single or sequential multiple derivative transitions for regenerative therapies.

## Author contributions

Conceptualization, M.L. and T.S.-S.; Methodology, M.L., S.M.; Validation, M.L., T.S.-S.; Analysis, M.L., T.S.-S; Writing, M.L., T.S.-S; Supervision, T.S.-S.; Funding Acquisition, T.S.-S.

## Acknowledgements

This work was supported by Wellcome Trust Senior Research Fellowship (215615/Z/19/Z) to T.S.S.; Radcliffe Department of Medicine Scholarship and MRC DTP Supplementary Funding to M.L. We express our gratitude to the Biomedical Services Aquatics team for the zebrafish maintenance. We thank the MRC Weatherall Institute of Molecular Medicine FACS and single cell facilities and Wolfson Imaging Centre. We also thank Steven Edwards and SciLifeLab Advanced Light Microscopy facility for their help with the Light Sheet Microscopy. Finally, we thank Life Science Editors for the help with manuscript editing and comments.

## Experimental Procedures

### Zebrafish husbandry

Animals were handled in accordance to procedures authorised by the UK Home Office in accordance with UK law (Animals [Scientific Procedures] Act 1986) and the recommendations in the Guide for the Care and Use of Laboratory Animals. All vertebrate animal work was performed at the facilities of Oxford University Biomedical Services. Adult fish were maintained as described previously (Westerfield, 2000). In brief, adult fish were exposed to 12 hour light – 12 hour dark cycle (8am to 10pm light; 10pm to 8am dark), kept in a closed recirculating system water at 27-28.5°C, fed 3-4 times a day, kept at 5 fish per 1L density. Embryos were staged as described previously (Kimmel et al., 1995). In brief, embryos were staged using a dissecting stereo-microscope: 1-2ss was recognised by observing first/second segment furrow; 5-6ss – counting 5/6 somites, apparent optical and Kupffer’s vesicles and prominent polster.

### Zebrafish lines

Genetrap lines, *Gt(foxd3-citrine)*^*ct110a*^ and *Gt(foxd3-mCherry)*^*ct110R*^, were generated by (Hochgreb-Hägele and Bronner, 2013). Enhancer reporter lines, *Tg(foxd3:enh6-EGFP)*^*ox109*^ and *Tg(foxd3:enh4-EGFP)*^*ox110*^, were generated cloning each enhancer from the genomic zebrafish DNA DNA using KAPA Long Range HotStart PCR kit (Kapa Biosystems) and cloned into the E1b:GFP:AC-DS vector (102417 Addgene (Chong-Morrison et al., 2018), linearised with NheI using the InFusion kit (638909, Takara Bio). Fertilised single-cell embryos were injected with 30 pg of plasmid DNA and 25 pg of *in vitro* transcribed Ac mRNA (using mMESSAGE mMACHINE SP6 Transcription Kit, AM1340 on a BamHI-linearised pAC-SP6 vector). F_0_ injected embryos were raised and screened for founders. Positive F_1_ embryos were raised for experiments.

Embryos at 5-6ss or larvae at 2dpf from *Tg(foxd3:enh6-EGFP)*^*ox109*^ or *Tg(foxd3:enh4-EGFP)*^*ox110*^ line were manually dechorionated if needed, anesthetised with tricaine and mounted in 1% low-melting agarose for live imaging. Embryos or larvae were imaged on a Zeiss780 LSM inverted confocal microscope equipped with Plan-APO Chromat 10×/0.45 NA WD=2.0 mm and LD C-APO Chromat 40×/1.1 NA WD=0.62 mm objectives or on a Zeiss780 LSM upright confocal microscope equipped with Plan-APO Chromat 10x/0.45 NA WD=2.0 mm objective. Images were processed using Fiji or Imaris image analysis software.

### Cell dissociation from zebrafish embryos and FAC-sorting

Selected embryos were dissociated with collagenase (20 mg/ml in 0.05% trypsin, 0.53 nM EDTA in 1X HBSS) in 300 ml volume at 30°C for 5-8 mins with intermittent pipetting to achieve a single cell suspension. The reaction was stopped by adding dissociation suspensions into 4 ml of Hanks buffer (10 mM Hepes (pH 8), 2.5 mg/ml of Bovine serum albumin (BSA) in 1X HBSS) followed by gentle trituration with a glass serological pipette. Cells were centrifuged at 500 xg for 10 min and re-suspended in 5 ml of Hanks buffer, passed through a 0.22 mm filter and centrifuged at 750 xg for 10 min, pelleted cells were resuspended in ~200-500 ml of Hanks buffer. Dead cells were stained with 7-AAD (A1310, ThermoFisher Scientific) and were excluded from collections. Fluorescent positive or negative cells were sorted and collected using BD FACSAria II SORP, BD FACSARIA III or BD FACSAria Fusion2.

### Single cell RNA-seq: RNA extraction, library preparation and sequencing

Embryos issued from incrosses of *Gt(foxd3-citrine)*^*ct110a*^ and *Gt(foxd3-mCherry)*^*ct110R*^ lines at 5-6ss were sorted for citrine fluorescence and dissociated to single cell suspension using the protocol described above. Individual citrine-positive (control) and citrine/mCherry (mutant) double positive cells were collected by FACS into two Eppendorf tubes, each containing 2 μl Hanks buffer. Sorted cells were loaded onto two Chromium 10X Genomics chips and the Chromium Single Cell 3’ Library and Gel Bead Kit v3 (Cat.No. PN-1000092) was used following manufacturer’s instructions. Libraries were quantified using Qubit (Cat. No.: Q32854) and KAPA library quantification kit (Cat.No.: KK4835). 4 nM 10X scRNA-seq libraries were sequenced on a NextSeq500 platform (Illumina) using NextSeq 500/550 High Output Kit v2.5 (150 Cycles) (Cat.No. 20024907) with 28 x 8 (i7) x 0 (i5) x 91 cycle number mode. Citrine/mCherry library was sequenced second time using the same materials and methods to acquire a sufficient number of reads.

Embryos issued from *Tg(foxd3:enh4-EGFP)*^*ox110*^ line at 90% epiboly and 1-2ss were sorted for EGFP fluorescence and processed for 10X Genomics scRNA-seq as above. Pooled 2 nM 10X scRNA-seq libraries were sequenced together on NextSeq500 platform using NextSeq 500/550 High Output Kit v2.5 (150 Cycles) with 28 × 8 (i7) × 0 (i5) × 91 cycle number mode.

### Single cell ATAC-seq: nuclei expulsion, tagmentation, library preparation and sequencing

Embryos issued from *Gt(foxd3-citrine)*^*ct110a*^ line at 5-6ss were sorted for citrine fluorescence and dissociated to single cell suspension using the protocol described above. Individual citrine-positive cells were collected by FACS into 2 μl Hanks buffer. Nuclei were isolated by lysing the cells with a hypotononic buffer for 5 min on ice following the 10X Genomics Low Cell Input Nuclei Isolation CG000169 protocol. Nuclei were transposed at 50°C for 1 hour, loaded onto the Chromium 10X Genomics chip and Chromium Single Cell ATAC Library & Gel Bead Kit (PN-1000111) was used following the CG000168 User guide. Library was quantified using Qubit and KAPA library quantification kit. 4 nM 10X scATAC-seq library was sequenced on a NextSeq500 platform (Illumina) using NextSeq 500/550 High Output Kit v2.5 (75 Cycles) (Cat.No. 20024906) with 34 × 34 × 8 (i7) × 16 (i5) cycle number mode.

### Enh6-EGFP+/− and Enh4-EGFP+/− cell bulk RNA extraction, library preparation and sequencing

Embryos issued from *Tg(foxd3:enh6-EGFP)*^*ox109*^ and *Tg(foxd3:enh4-EGFP)*^*ox110*^ lines at 5-6ss were sorted for citrine fluorescence and dissociated to single cell suspension using the protocol described above. FAC sorted cells (5000 cells for each sample: Enh6-EGFP+ (in triplicates), Enh6-EGFP-(in duplicates), Enh4-EGFP+ (in triplicates) and Enh4-EGFP-(in duplicates)) were washed with PBS and stored in lysis buffer (RNAqueous-Micro Total RNA Isolation Kit, Ambion, AM1931) at −80°C. RNA was extracted using Ambion RNAqueous Micro Total RNA isolation kit and checked on TapeStation using Agilent High Sensitivity RNA Tape (Cat.No.:5067-5579) and buffer (Cat.No.:5067-5580). cDNA was prepared using Takara Clontech SMART-Seq v4 Ultra Low Input RNA Kit for Sequencing (Cat.No.:634890) using 8 cycles of amplification and purified using Agencourt AMPure XP beads (Beckman Coulter, A63880). 1 ng of each sample cDNA was used for library preparation. Sequencing libraries were prepared using Illumina Nextera XT library preparation kit (FC-131-1096) and purified using Agencourt AMPure XP beads. The library quality was checked using Agilent High Sensitivity TapeStation D1000 ScreenTape (Cat. No.: 5067-5584) and Reagents (Cat. No.: 5067-5585). All libraries were pooled into a single pool, quantified using KAPA library quantification kit and sequenced using NextSeq 500/550 High Output Kit v2 (75 cycles, paired-end 40 bp run) on NextSeq 500 sequencing system.

### Enh6-EGFP+/− and Enh4-EGFP+/− cell bulk ATAC-seq library preparation and sequencing

Embryos issued from *Tg(foxd3:enh6-EGFP)*^*ox109*^ and *Tg(foxd3:enh4-EGFP)*^*ox110*^ lines at 5-6ss were sorted for citrine fluorescence and dissociated to single cell suspension using the protocol described above. FAC sorted cells (15,000 - 25,000 for each sample in triplicates: Enh6-EGFP+, Enh6-EGFP-, Enh4-EGFP+ and Enh4-EGFP-) were lysed (10 mM Tris-HCl, pH7.4, 10 mM NaCl, 3 mM MgCl2, 0.1% Igepal) and tagmented using Nextera DNA kit (Illumina FC-121-1030) (tagmentation −0.7 μl of Tn5 enzyme in 10 μl reaction) for 30 min at 37°C. Reactions were quenched with 50 mM EDTA for 30 min at 50°C. Tagmented fragments were purified using MinElute PCR Purification Kit (Cat No.: 28004). Tagmented DNA was amplified using NEB Next High-Fidelity 2X PCR Master Mix (Cat. No.: M0541S) for 12-13 cycles and purified using Agencourt AMPure XP beads. Tagmentation efficiency and final libraries were assessed using Agilent High Sensitivity TapeStation D1000 ScreenTape and Reagents. All libraries were pooled into a single pool, quantified using KAPA library quantification kit and sequenced using NextSeq 500/550 High Output Kit v2 (75 cycles, paired-end 40 bp run) on NextSeq 500 sequencing system.

### In situ Hybridisation Chain Reaction (HCR)

HCR kit (v3, Molecular Instruments (Choi et al., 2018)) amplification and hybridisation buffers and wash buffers, fluorescently labelled hairpins and commercial or custom-made DNA probe sets was used for each multiplexed target mRNA *in situ* visualisations. Zebrafish embryos were dechorionated at desired stages and fixed in 4% PFA for 24 hours at 4ºC. HCR was performed following the ‘In situ HCR v3.0 protocol for whole-mount zebrafish larvae’ protocol. Ethanol instead of methanol was used for embryo dehydration/rehydration. Embryos were mounted in 1% low melting-agarose and imaged on Zeiss780/880 LSM inverted confocal microscopes equipped with Plan-APO Chromat 10x/0.45 NA WD=2.0 mm and LD C-APO Chromat 40x/1.1 NA WD=0.62 mm objectives; on a Zeiss780 LSM upright confocal microscope equipped with Plan-APO Chromat 10x/0.45 NA WD=2.0 mm objective or on a Zeiss Lighsheet Z.1 microscope equipped with 10x/0.5 NA water dipping detection objective. Images were processed using Fiji or Imaris image analysis software.

### Statistical analysis and bioinformatics data processing

#### 5-6ss foxd3 control and mutant scRNA-seq analysis

Single-cell RNA-seq raw base call files were demultiplexed using Cellranger (v3.1.0) (Zheng et al., 2017), mapped to the Danio rerio.GRCz11 (danRer11) transcriptome, that had Citrine and mCherry transcripts added to the FASTA file, and compiled using the Cellranger ‘mkref’ function using Danio rerio.GRCz11.99.chr.filtered.gtf and Danio rerio.GRCz11.dna.primary assembly.fa annotations. 5,212 estimated number of foxd3-ctrine (control) cells were obtained with mean reads per cell of 75,607, median genes per cell of 3,854, median UMI counts per cell of 22,636 and total number of genes detected was 21,465. 97.4% of reads had valid barcodes with a Q30 of 97% and 89.9% of the reads mapped confidently to the zebrafish genome. 3,615 estimated number of foxd3-ctrine/mCherry (mutant) cells were obtained with mean reads per cell of 32,281, median genes per cell of 2,828, median UMI counts per cell of 12,787 and total number of genes detected was 20,240. 93.1% of reads had valid barcodes with a Q30 of 98% and 89.2% of the reads mapped confidently to the zebrafish genome. Downstream analysis was carried out using Seurat package (v3.0.0) in R (Butler et al., 2018). Matrices were filtered to remove cells with fewer than 500 genes and more than 5000 genes detected, and with more than 8.5% of mitochondrial genes detected, resulting in 3,684 remaining control and 3,420 mutant cells used for analysis. Expression values for the total UMI counts per cell were normalised and permutation test carried to determine the significant principal components and dimensions in the data before performing linear dimensional reduction (resolution = 0.5, dims = 1:20). Clusters were visualised on Umap plots. Control Trunk NC (NC-1), Mixed NC, NMps/NMp mesoderm, Paraxial mesoderm or mutant Trunk NC (NC-1), Mixed NC, NMps, NMp-neural, Paraxial mesoderm clusters were selected for further sub-clustering using resolution = 0.5 (for control), resolution = 0.4 (for mutant) and dims = 1:12 parameters. Clusters were annotated manually based on enriched marker genes. Statistically significant differential gene expression between clusters was identified using Wilcoxon rank sum test.

The ‘Mixed NC’ cluster exhibited lower UMI count in comparison to other clusters (fig. S1 C). To make sure these cells did not cluster together because of that and also to investigate whether the lower UMI count is biologically meaningful, we have again filtered all the initially sequenced cells with, this time, 350-1500 UMIs and with <5% expressed mitochondrial gene content. We also regressed mitochondrial/ribosomal transcript and UMI counts during the filtered cell clustering, consequently uncovering 4 distinctive clusters (fig. S1 D-E). While Cluster 3 cells were mainly expressing various keratins, Clusters 0-2 exhibited somewhat mixed NMp-NC signatures. Cluster 1 was mostly enriched for the NMp markers (*cdx4*, *tbx16*, *msgn1*, *sall4*), Cluster 2 – NMp-neural markers (*alcama*, *nova2*), but also NC genes (*zic2b*, *tfap2a*, *marcksl1b*, *foxd3*) (fig. S1 F). Finally, Cluster 0 cells were enriched for the same NC genes, but also NMp and NMp-neural related genes (*tbx16*, *cdx4*, *sall4*, *msgn1*, *nova2*, *alcama*) (fig. S1 F).

3,684 control and 3,420 mutant cells were integrated into a single reference by finding integration anchor points between two data sets using the Seurat pipeline (Butler et al., 2018). Integrated Seurat object was clustered using resolution = 0.5 and dims = 1:20 parameters. Clusters were annotated manually based on enriched marker genes. Statistically significant differential gene expression between specified mutant and control clusters was identified using Wilcoxon rank sum test.

#### 90% epiboly/1-2ss Enh4-EGFP+ single cell analysis

Single-cell RNA-seq raw base call files were demultiplexed using Cellranger (v3.1.0) (Zheng et al., 2017), mapped to the Danio rerio.GRCz11 (danRer11) transcriptome and compiled using the Cellranger ‘mkref’ function. 2,671 estimated number of 90% epiboly cells were obtained with mean reads per cell of 91,291, median genes per cell of 2,954, median UMI counts per cell of 14,958 and total number of genes detected was 19,731. 94.9% of reads had valid barcodes with a Q30 of 92.3% and 70.6% of the reads mapped confidently to the zebrafish genome. 5,425 estimated number of 1-2ss cells were obtained with mean reads per cell of 62,622, median genes per cell of 2,654, median UMI counts per cell of 13,403 and total number of genes detected was 20,672. 95.1% of reads had valid barcodes with a Q30 of 92.5% and 75.9% of the reads mapped confidently to the zebrafish genome. Downstream analysis was carried out using Seurat package (v3.0.0) in R (Butler et al., 2018). Both samples sequenced together were merged using ‘merge’ Seurat function. Combined matrices were filtered to remove cells with fewer than 500 genes and more than 5500 genes detected, and with more than 5% of mitochondrial and 40% ribosomal genes detected. Expression values for the total UMI counts per cell were normalised and permutation test carried to determine the significant principal components and dimensions in the data before performing linear dimensional reduction (resolution = 0.5, dims = 1:15). UMI, mitochondrial and ribosomal gene counts were regressed out during clustering. One outlier cluster with low UMI counts and not specifically enriched genes was removed from further analysis. Clusters of final 5,123 cells were visualised on Umap plots and annotated manually based on enriched marker genes.

#### RNA velocity trajectory and latent time analysis

The analysis was performed using the scVelo dynamical modelling pipeline (Bergen et al., 2020) in python. For this, loom files containing spliced/unspliced transcript expression matrices were generated using velocyto.py pipeline (La Manno et al., 2018). “Loom cell” names were renamed to match Seurat object cell names and only Seurat-filtered cells were selected for trajectory analysis. Seurat generated UMAP coordinates, clusters and cluster colours were added to the filtered “loom cells”. Another dimensionality reduction package (scanpy.external.pl.phate), Phate, was used with the default parameters (Moon et al., 2019). Pseudotime package Slingshot (Street et al., 2018) was used with stretch = 0 and starting with cluster NMp-2, NMp-1 or NMp-neural giving the same results.

#### scATAC-seq from 5-6ss foxd3-citrine cells analysis

Single cell ATAC-seq raw base call files were demultiplexed using cellranger-atac/1.1.0 and mapped to the Danio rerio.GRCz11 (danRer11) genome generated with ‘mkref’ function using Danio rerio.GRCz11.99.chr.filtered.gtf and Danio rerio.GRCz11.dna.primary assembly.fa annotations. 1,879 estimated number of foxd3-ctrine cells were obtained with 43,010 median fragments per cell, 97.6% fraction of read pairs with a valid barcode, 89.9% Q30 bases in barcode, 67.3% fraction of total read pairs mapped confidently to genome (>30 mapq). Downstream analysis was carried out using Signac (1.0.0) and Seurat in R (Stuart et al., 2020). Cells with 2000 – 100,000 peak region fragments, more than 15% of reads in peaks, nucleosomal signal smaller than 10 and transcriptional start site enrichment higher than 2 were kept for downstream analysis resulting in 946 remaining cells. Term frequency-inverse document frequency (TF-IDF) normalization and latent semantic indexing (lsi) dimension reduction method was used to cluster cells (dims = 2:30, res = 1.2). Clusters were visualised on Umap plots. Gene activity matrix to visualise selected gene accessibility on a UMAP plot was built by assessing the chromatin accessibility intersecting each gene promoter and gene body. scATAC clusters were labelled by using foxd3-citrine scRNA-seq clusters as reference to find anchors and transfer labels by employing canonical correlation analysis (CCA) and lsi for integrated object dimension reduction. Cells with the prediction score higher than 0.3 were subset.

Foxd3-citrine scRNA-seq (clusters: NC-1, NC-3, Cranial NC, NMps/NMp-mesoderm, ‘Mixed NC’, Noto-pr. and Paraxial mesoderm) and scATAC-seq cell clusters were co-embedded into a single reference by performing imputation to impute scRNA-seq matrix for each scATAC-seq cell, using lsi weight reduction and ‘merge’ function, scaling data to variable features, and using dims = 10 for clustering and UMAP visualisation.

#### Enh6-EGFP+/− and Enh4-EGFP+/− cell bulk RNA-seq bioinformatics processing

Reads were trimmed to remove low quality bases using trim galore/0.4.1 (Babraham). Read quality was evaluated using FastQC (Babraham). Mapping to GRCz10/danRer10 assembly of the zebrafish genome downloaded from UCSC Genome Browser was performed using rna-star/2.4.2a (Dobin et al., 2013). Read counts were obtained using subread Feature-Counts (subread/1.6.2) (Liao et al., 2014) with standard parameters using a gene model gtf derived from Ensembl annotation downloaded from UCSC genome browser. Differential Expression analysis (Enh6+ vs Enh6-, Enh4+ vs Enh4- and Enh6+ vs Enh4+) was carried out using DESeq2 (1.24.0) (Love et al., 2014). Gene Ontology Biological Terms were acquired using PANTHER (Mi et al., 2019) Danio Rerio terms (used parameters: transcripts expressed at basemean>50, with differential expression at LogFC>2 or LogFC<-2 between compared groups were used for GO biological process complete, Fisher test with false discovery rate (FDR) correction. GO terms were visualised as a bubble plot using ggplot2 (Wickham, 2016) package in R.

#### Enh6-EGFP+/− and Enh4-EGFP+/− cell bulk ATAC-seq bioinformatics processing

Reads were trimmed to remove low quality bases using trim galore/0.4.1 (Babraham). Read quality was evaluated using FastQC (Babraham). To visualise foxd3 locus for the ATAC-seq results, reads mapping to the enhancer reporter plasmid were excluded by: first mapping all acquired reads (using bowtie/1.1.2.) to a plasmid sequence carrying 150bp of a cloned Enh6 or Enh4 sequence from each side of a plasmid, and then taking all the unmapped reads for the downstream mapping to the danRer10 genome using bowtie/1.1.2. Duplicates were removed using Picard tools MarkDuplicates (picard-tools/1.83). Sample reads were equalised to the same number by random down-sampling using samtools-1.1 (to 24,661,382 reads, except for one of Ehh4-samples that had 11,457,938 reads). BAM files were sorted by name and paired end bed files were obtained using bedtools (v.2.15.0) bamtobed -bedpe. Reads that were not properly paired were discarded and paired reads were displaced by +4 bp and -5 bp for transposition correction. Bigwig files were generated using an enhanced Perl script (courtesy of Prof Jim Hughes) and visualised on the UCSC Genome Browser.

#### JASPAR and HOCOMOCO transcription factor binding site (TFBS) analysis

TFBS analysis on the Enh6 or the Enh4 DNA sequences, which were cloned into the E1b-GFP-AC-DS vector used for the transgenesis, was performed using both, HOCOMOCO (http://hocomoco11.autosome.ru) and JASPAR (7th release, http://jaspar.genereg.net/) databases (Khan et al., 2018; Kulakovskiy et al., 2018). For the JASPAR analysis, Vertebrata taxonomic group database was used, set to 87% relative profile score threshold. Candidate TFs were then selected based on identified multiple predicted binding sites for the same TF family for either Enh4 or Enh6 sequences. For the HOCOMOCO analysis, mouse mononucleotide P-value 0.0001 TFBS model motif in Homer format was scanned and annotated using Homer (v.4.8) annotatePeaks.pl (Heinz et al., 2010). Candidate TFs were then selected based on identified multiple predicted binding sites for the same TF family for either Enh4 or Enh6 sequences. For final TFBS plotting: to highlight the most enriched and also differential TFBSs underlying Enh6 and Enh4 DNA sequences, we have prioritised and plotted those uncovered TFBSs that were found: (a) at least 3 times within either Enh6 or Enh4 and (b) each uncovered TF had at least two more sites between Enh6 and Enh4 or Enh4 and Enh6 sequences.

#### NMp differentiation at 5-6ss gene regulatory network (GRN) building

In order to access all the scATAC peaks specifically found in the NMp cells, we have employed the Loupe Cell browser for scATAC data, that also distinguished the equivalent NMp-cell population exhibiting NMp genespecific accessibility profiles. We extracted the NMp-specific cluster peaks (cl.2 from the k-mean clustering, fig. S7, b-f) from the dataset, annotated them to the nearest genes, and zoomed in on the scRNA-seq genes previously identified to be enriched in either NMps, NMp-neural cells or MPCs (NMp-related cells) (fig. S6 d). DanRer11 genome assembly coordinates were converted to DanRer10 assembly coordinates using LiftOver tool from UCSC Genome Browser. We then annotated pre-selected (expressed TFs in the NMp-related cells) vertebrate TF motifs (P<0.0001) from the Hocomoco v11 database (Kulakovskiy et al., 2018) across these three groups using Homer (v.4.8) annotatePeaks.pl (-size 500, in-built danRer10 genome), the acquired motif annotation lists were converted to a matrix of a motif occurrence numbers for each peak using a custom R script. For network generation a single TF motif for each gene was used. The circuits of the selected core factors were built using BioTapestry tool (Longabaugh et al., 2009).

