## Supplementary figures for "Neuromesodermal progenitor origin of trunk neural crest *in vivo*"

---

**Supplemental Material**

---

[1]University of Oxford, MRC Weatherall Institute for Molecular Medicine, Radcliffe Department of Medicine, Oxford OX3 9DS, UK
[2]Current address: Department of Cell and Molecular Biology, Karolinska Institutet, SE-171 77 Stockholm, Sweden

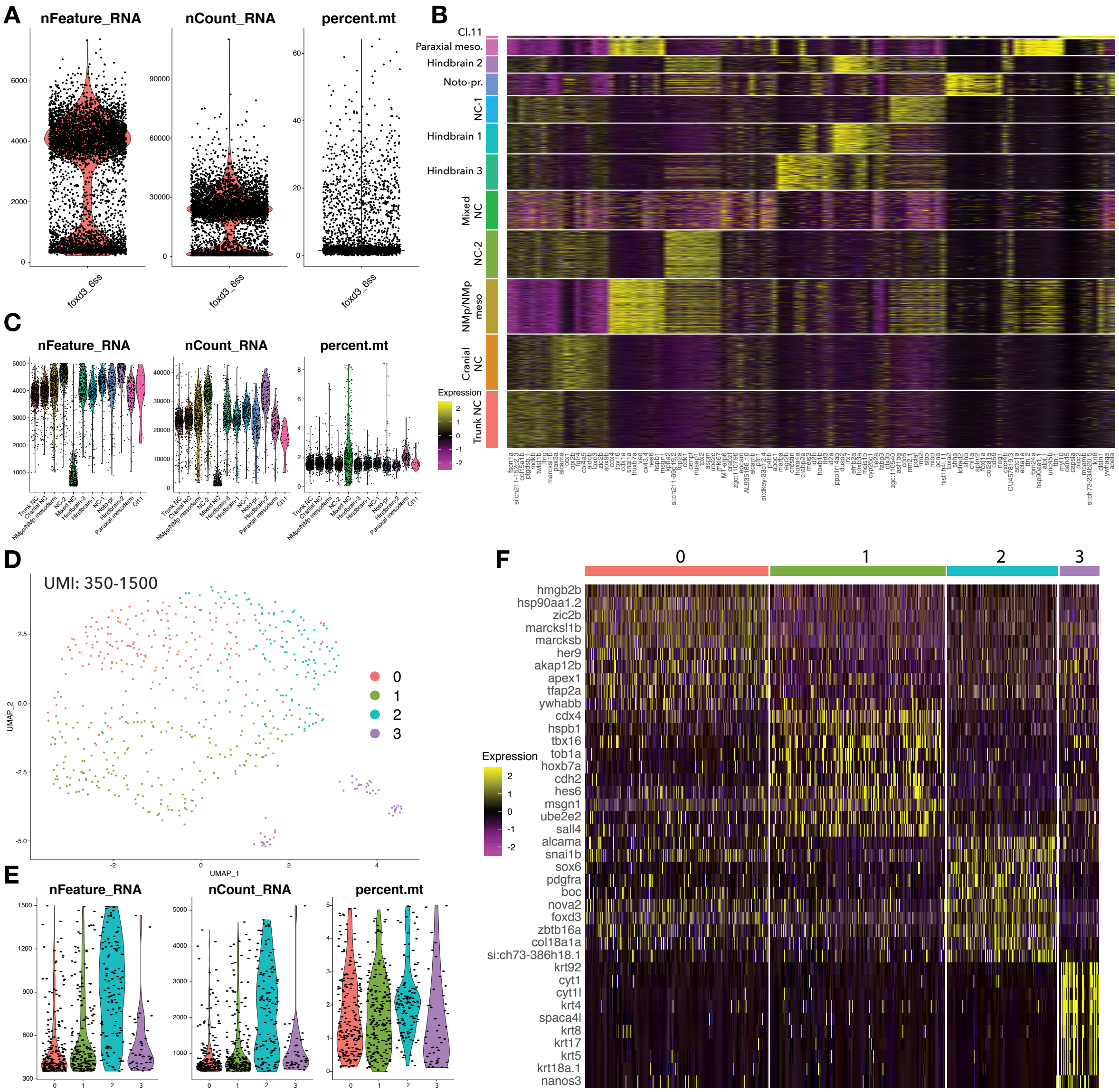
A
nFeature_RNA
nCount_RNA
percent.mt
foxd3_6ss
B
Cl.11
Paraxial meso.
Hindbrain 2
Noto-pr.
NC-1
Hindbrain 1
Hindbrain 3
Mixed NC
NC-2
NMp/NMp meso
Cranial NC
Trunk NC
Expression
C
nFeature_RNA
nCount_RNA
percent.mt
D
UMI: 350-1500
UMAP_1
UMAP_2
0
1
2
3
E
nFeature_RNA
nCount_RNA
percent.mt
F
0
1
2
3
Expression
hmgb2b
hsp90aa1.2
zic2b
marcksl1b
marcksb
her9
akap12b
apex1
tfap2a
ywhabb
cdx4
hspb1
tbx16
tob1a
hoxb7a
cdh2
hes6
msgn1
ube2e2
sall4
alcama
snai1b
sox6
pdgfra
boc
nova2
foxd3
zbtb16a
col18a1a
si:ch73-386h18.1
krt92
cyt1
cyt1l
krt4
spaca4l
krt8
krt17
krt5
krt18a.1
nanos3

S1

**Figure S1**

**Foxd3+(citrine+) cell scRNA-seq data processing and clustering. (A)** Violin plots of sequenced/unfiltered cell quality metrics. **(B)** Top 10 markers enriched in each foxd3+ cell clusters. **(C)** Violin plots of quality metrics grouped by cluster. **(D)** Uniform Manifold Approximation and Projection (UMAP) embedding of foxd3+ cells filtered for 350-1500 unique molecular identifier (UMI) counts. **(E)** Violin plots of quality metrics grouped by **(D)** cluster. **(F)** Top 10 markers enriched in each **(D)** cell cluster.

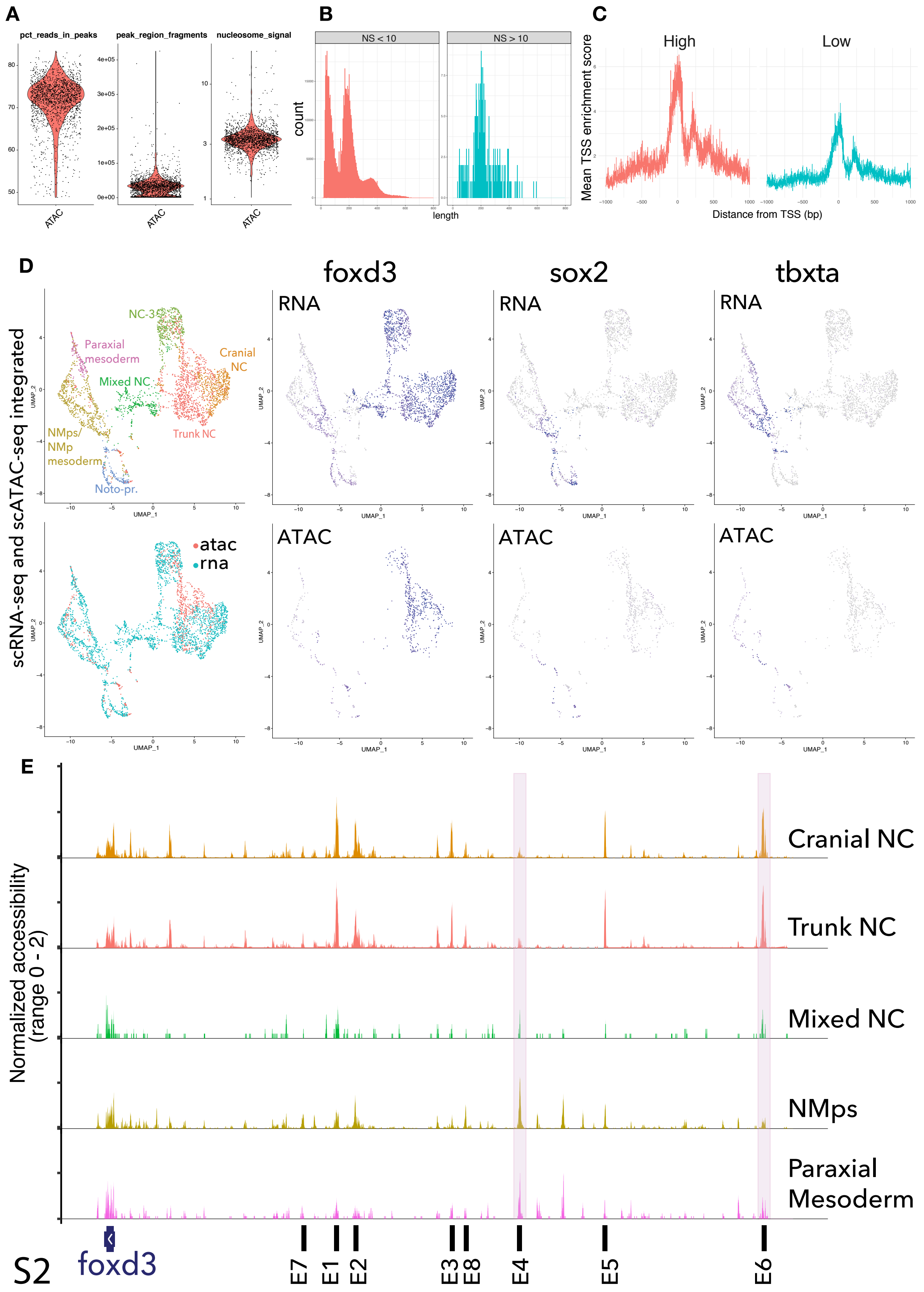

A
pct_reads_in_peaks
peak_region_fragments
nucleosome_signal
ATAC
B
NS < 10
NS > 10
count
length
C
High
Low
Mean TSS enrichment score
Distance from TSS (bp)
D
foxd3
sox2
tbxta
RNA
ATAC
scRNA-seq and scATAC-seq integrated
NC-3
Paraxial mesoderm
Cranial NC
Mixed NC
Trunk NC
NMps/ NMp mesoderm
Noto-pr.
atac
rna
UMAP_1
UMAP_2
E
Normalized accessibility (range 0 - 2)
Cranial NC
Trunk NC
Mixed NC
NMps
Paraxial Mesoderm
foxd3
E7
E1
E2
E3
E8
E4
E5
E6

S2

**Figure S2**

**Foxd3+(citrine+) cell scATAC-seq data processing and integration with foxd3+ (citrine+) cell scRNA-seq.** **(A)** Violin plots of sequenced/unfiltered single cell ATAC quality metrics. **(B)** Quality metrics of unfiltered cells for fragment length periodicity grouped by high or low nucleosomal signal (averaged from 10Mb region on chr1) and **(C)** high or low transcription-start site (TSS) enrichment score (average of 2000 TSSs). Cells with 100000 $<$ peak region fragments $>$ 2000 & percentage of reads in peaks $>$ 15 & nucleosome_signal $<$ 10 & TSS.enrichment $>$ 2 were filtered for downstream analysis. **(D)** UMAP embedding for foxd3+ cell scRNA-seq and scATAC-seq data integration into a single reference; cell integration visualisation; and foxd3, sox2 and tbxta gene expression and accessibility visualisation in clusters. **(E)** *foxd3* genomic locus accessibility across selected scATAC-seq clusters; enhancer 4 (Enh4) and enhancer 6 (Enh6) are highlighted in pink boxes.

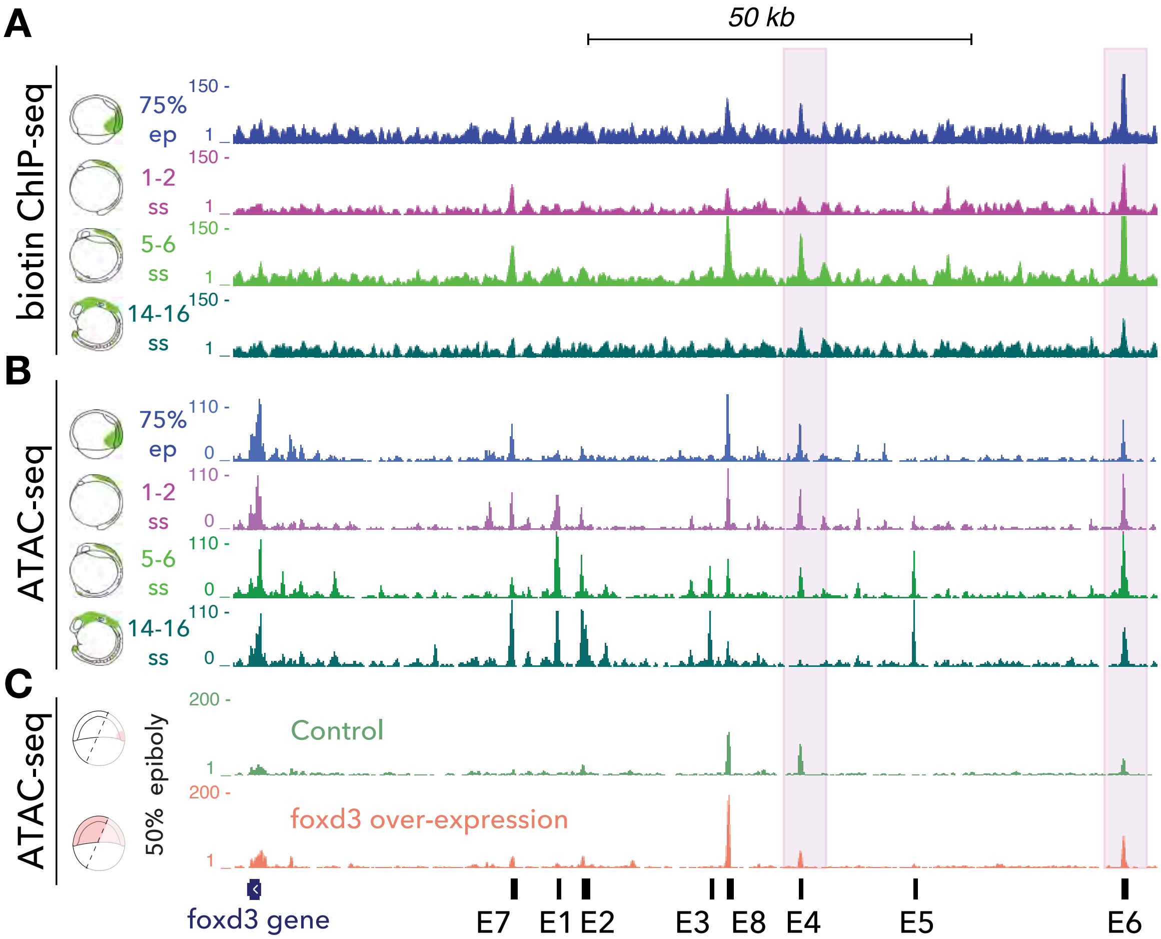

A
50 kb
biotin ChIP-seq
75% ep
1-2 ss
5-6 ss
14-16 ss
150 -
1
B
ATAC-seq
110 -
0
C
ATAC-seq
50% epiboly
200 -
Control
foxd3 over-expression
foxd3 gene
E7
E1
E2
E3
E8
E4
E5
E6


S3

### Figure S3

***Foxd3* gene locus and auto-regulated *cis*-regulatory elements. (A)** Genome browser visualisation depicting region upstream of the *foxd3* transcription start site, containing putative *foxd3* *cis*-regulatory elements. Foxd3 Biotin ChIP-seq at 75% epiboly (blue), 1-2 somite (purple), 5-6 somite (light green) and 14 somite (dark green) stages. **(B)** ATAC-seq from foxd3+(citrine+) cells at 75% epiboly (blue), 1-2 somite (purple), 5-6 somite (light green) and 14-16 somite (dark green) stages. **(C)** Upon *foxd3* mRNA injection into single cell stage embryos, embryos were collected for ATAC-seq at 50% epiboly stage. Embryos were dissected (dashed lines) to only collect "foxd3-naïve" cells that do not normally express *foxd3*. Native and ectopic *foxd3* expression is illustrated in dark pink and lighter pink, respectively. Green and light orange ATAC-seq tracks represent genome accessibility from control and *foxd3* overexpressing embryonic cells from 50% stage embryos, respectively.

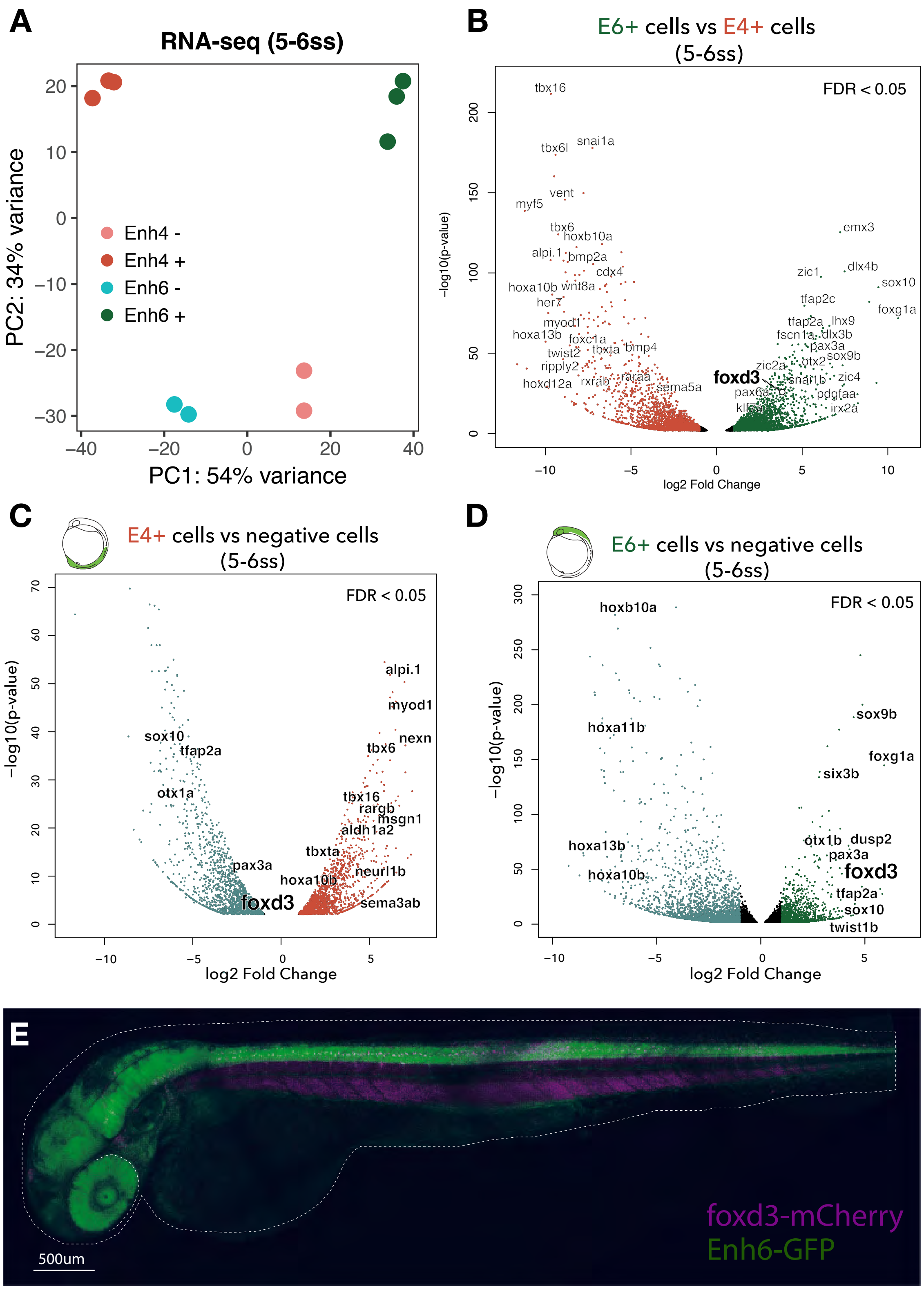
A
RNA-seq (5-6ss)
PC2: 34% variance
PC1: 54% variance
Enh4 -
Enh4 +
Enh6 -
Enh6 +
B
E6+ cells vs E4+ cells
(5-6ss)
FDR < 0.05
-log10(p-value)
log2 Fold Change
tbx16
snai1a
tbx6l
vent
myf5
tbx6
hoxb10a
alpi.1
bmp2a
cdx4
hoxa10b
wnt8a
her7
myod1
hoxa13b
foxc1a
twist2
tbxta
bmp4
ripply2
rxrab
raraa
hoxd12a
sema5a
foxd3
emx3
dlx4b
zic1
sox10
tfap2c
foxg1a
tfap2a
lhx9
fscn1a
dlx3b
pax3a
otx2
sox9b
zic2a
snai1b
zic4
pax6a
pdgfaa
irx2a
C
E4+ cells vs negative cells
(5-6ss)
FDR < 0.05
−log10(p-value)
log2 Fold Change
alpi.1
myod1
nexn
tbx6
sox10
tfap2a
otx1a
tbx16
rargb
msgn1
aldh1a2
tbxta
pax3a
hoxa10b
neurl1b
foxd3
sema3ab
D
E6+ cells vs negative cells
(5-6ss)
FDR < 0.05
−log10(p-value)
log2 Fold Change
hoxb10a
sox9b
hoxa11b
foxg1a
six3b
hoxa13b
otx1b
dusp2
pax3a
hoxa10b
foxd3
tfap2a
sox10
twist1b
E
foxd3-mCherry
Enh6-GFP
500um

S4

### Figure S4

**Profiling Enh4+ *Tg(foxd3:enh4-EGFP)$^{ox110}$* and Enh6+ *Tg(foxd3:enh6-EGFP)$^{ox109}$* cells by bulk RNA-seq at 5-6 somite stage (ss). (A)** Triplicate samples from either *Tg(foxd3:enh4-EGFP)$^{ox110}$* or *Tg(foxd3:enh6-EGFP)$^{ox109}$* embryos were dissociated and EGFP+ fluorescent embryonic cells were isolate using Fluorescence-activated cell sorting (FACS) method. Corresponding negative (EGFP-) cells were also collected in duplicates as negative control samples. RNA-seq libraries for all samples were generated and sequenced. PCA analysis shows four distinctive groups based on differential gene expression (DeSeq2): Enh6+, Enh6-, Enh4+ and Enh4- groups. **(B)** Volcano plot depicting differential expression analysis performed between Enh6+ and Enh4+ cells. Only those genes that were found to be bellow false discovery rate (FDR) of 0.05 were taken into account. Dots in black correspond to genes between (0 > log2fold < 1); in dark green dots - genes at (log2fold > 1) change, hence enriched in the Enh6+ cells when compared to Enh4+ cells; in red dots - genes at (log2fold < -1) change, hence enriched in the Enh4+ cells when compared to Enh6+ cells. **(C)** Volcano plot depicting differentially expressed genes between Enh4+ (in red) and Enh4- cells (in teal green) (FDR<0.05). **(D)** Volcano plot depicting differentially expressed genes between Enh6+ (in dark green) and Enh6- cells (in teal green) (FDR<0.05). **(E)** Lateral view of 2 days post fertilisation (dpf) zebrafish larva (*Gt(foxd3-mCherry)$^{ct110aR}$* transgenic line crossed with *Tg(foxd3:enh6-EGFP)$^{ox109}$*) expressing foxd3-mcherry (coloured in magenta) and Enh4-GFP (in green).

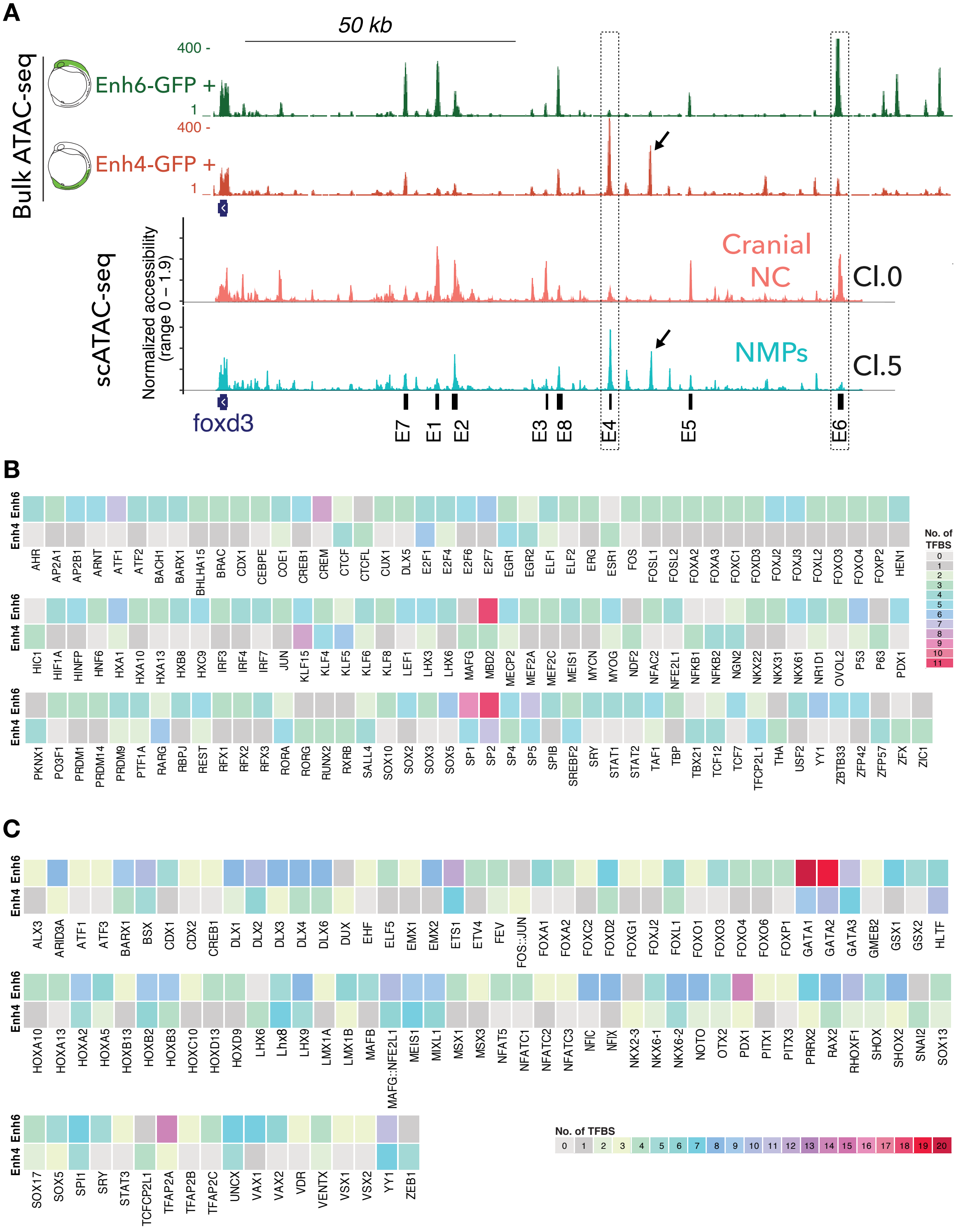
A
50 kb
Bulk ATAC-seq
Enh6-GFP +
Enh4-GFP +
400 -
1 _
scATAC-seq
Normalized accessibility (range 0 – 1.9)
Cranial NC
Cl.0
NMPs
Cl.5
foxd3
E7 E1 E2 E3 E8 E4 E5 E6
B
Enh6
Enh4
AHR AP2A1 AP2B1 ARNT ATF1 ATF2 BACH1 BARX1 BHLHA15 BRAC CDX1 CEBPE COE1 CREB1 CREM CTCF CTCFL CUX1 DLX5 E2F1 E2F4 E2F6 E2F7 EGR1 EGR2 ELF1 ELF2 ERG ESR1 FOS FOSL1 FOSL2 FOXA2 FOXA3 FOXC1 FOXD3 FOXJ2 FOXJ3 FOXL2 FOXO3 FOXO4 FOXP2 HEN1
HIC1 HIF1A HINFP HNF6 HXA1 HXA10 HXA13 HXB8 HXC9 IRF3 IRF4 IRF7 JUN KLF15 KLF4 KLF5 KLF6 KLF8 LEF1 LHX3 LHX6 MAFG MBD2 MECP2 MEF2A MEF2C MEIS1 MYCN MYOG NDF2 NFAC2 NFE2L1 NFKB1 NFKB2 NGN2 NKX22 NKX31 NKX61 NR1D1 OVOL2 P53 P63 PDX1
PKNX1 PO3F1 PRDM1 PRDM14 PRDM9 PTF1A RARG RBPJ REST RFX1 RFX2 RFX3 RORA RORG RUNX2 RXRB SALL4 SOX10 SOX2 SOX3 SOX5 SP1 SP2 SP4 SP5 SPIB SREBF2 SRY STAT1 STAT2 TAF1 TBP TBX21 TCF12 TCF7 TFCP2L1 THA USF2 YY1 ZBTB33 ZFP42 ZFP57 ZFX ZIC1
No. of TFBS
0 1 2 3 4 5 6 7 8 9 10 11
C
ALX3 ARID3A ATF1 ATF3 BARX1 BSX CDX1 CDX2 CREB1 DLX1 DLX2 DLX3 DLX4 DLX6 DUX EHF ELF5 EMX1 EMX2 ETS1 ETV4 FEV FOS::JUN FOXA1 FOXA2 FOXC2 FOXD2 FOXG1 FOXJ2 FOXL1 FOXO1 FOXO3 FOXO4 FOXO6 FOXP1 GATA1 GATA2 GATA3 GMEB2 GSX1 GSX2 HLTF
HOXA10 HOXA13 HOXA2 HOXA5 HOXB13 HOXB2 HOXB3 HOXC10 HOXD13 HOXD9 LHX6 Lhx8 LHX9 LMX1A LMX1B MAFB MAFG::NFE2L1 MEIS1 MIXL1 MSX1 MSX3 NFATC5 NFATC1 NFATC2 NFATC3 NFIC NFIX NKX2-3 NKX6-1 NKX6-2 NOTO OTX2 PDX1 PITX1 PITX3 PRRX2 RAX2 RHOXF1 SHOX SHOX2 SNAI2 SOX13
SOX17 SOX5 SPI1 SRY STAT3 TCFCP2L1 TFAP2A TFAP2B TFAP2C UNCX VAX1 VAX2 VDR VENTX VSX1 VSX2 YY1 ZEB1
No. of TFBS
0 1 2 3 4 5 6 7 8 9 10 11 12 13 14 15 16 17 18 19 20

S5

**Figure S5**

**Profiling Enh4 and Enh6 accessibility and enriched transcription factor binding sites (TFBSs (A)** Surveying and comparing open chromatin regions within the foxd3 locus between Enh6+ and Enh4+ cells and scATAC-seq Cranial NC and NMp clusters. Top two tracks - bulk ATAC-seq tracks from cells expressing Enh6 (in green) and Enh4 (in red) reporters. ATAC-seq reads were corrected for the transgenic Enh4 or Enh6 plasmid mapping. Bottom two pseudo-bulk tracks belong to scATAC-seq Cl.0 (Cranial NC) and Cl5. (NMp) clusters. Black arrows highlight an NMp/Enh4+ cell population-specific open chromatin region, which was not observed from our previous NC-specific ATAC-seq data sets. **(B)** TFBS analysis underlying Enh6 and Enh4 DNA sequences using JASPAR database. Predicted TFs were listed alphabetically and a number of each predicted TFBS (from 0 to 20) was illustrated using different colours. **(C)** TFBS analysis underlying Enh6 and Enh4 DNA sequences using HOCOMOCO database. Predicted TFs were listed alphabetically and a number of each predicted TFBS (from 0 to 11) was illustrated using different colours.

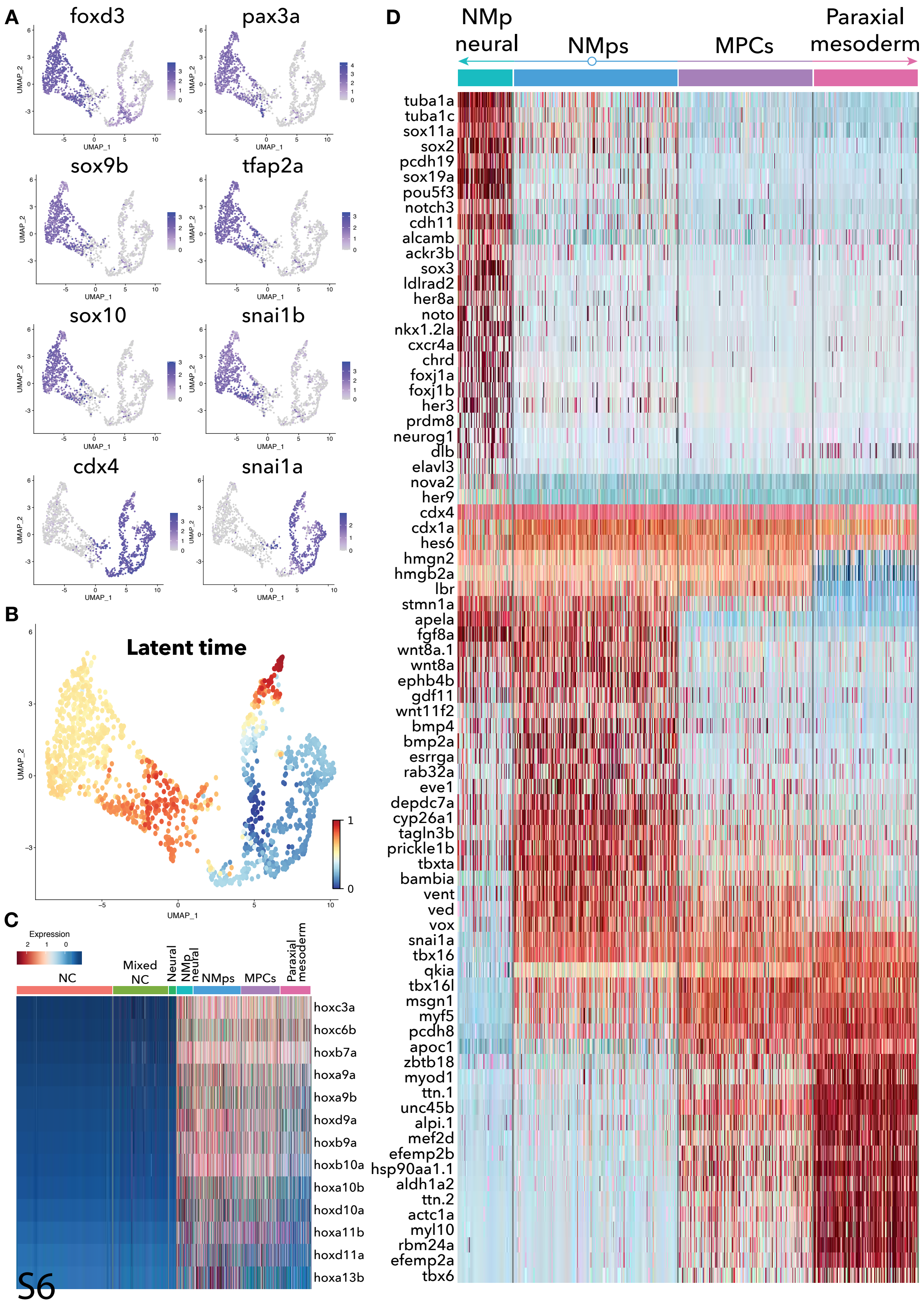
A
foxd3
pax3a
sox9b
tfap2a
sox10
snai1b
cdx4
snai1a
UMAP_1
UMAP_2
B
Latent time
C
Expression
NC
Mixed NC
Neural
NMp neural
NMps
MPCs
Paraxial mesoderm
hoxc3a
hoxc6b
hoxb7a
hoxa9a
hoxa9b
hoxd9a
hoxb9a
hoxb10a
hoxa10b
hoxd10a
hoxa11b
hoxd11a
hoxa13b
S6
D
NMp neural
NMps
MPCs
Paraxial mesoderm
tuba1a
tuba1c
sox11a
sox2
pcdh19
sox19a
pou5f3
notch3
cdh11
alcamb
ackr3b
sox3
ldlrad2
her8a
noto
nkx1.2la
cxcr4a
chrd
foxj1a
foxj1b
her3
prdm8
neurog1
dlb
elavl3
nova2
her9
cdx4
cdx1a
hes6
hmgn2
hmgb2a
lbr
stmn1a
apela
fgf8a
wnt8a.1
wnt8a
ephb4b
gdf11
wnt11f2
bmp4
bmp2a
esrrga
rab32a
eve1
depdc7a
cyp26a1
tagln3b
prickle1b
tbxta
bambia
vent
ved
vox
snai1a
tbx16
qkia
tbx16l
msgn1
myf5
pcdh8
apoc1
zbtb18
myod1
ttn.1
unc45b
alpi.1
mef2d
efemp2b
hsp90aa1.1
aldh1a2
ttn.2
actc1a
myl10
rbm24a
efemp2a
tbx6

### Figure S6

**Gene dynamics underlying neuromesodermal progenitor (NMp) differentiation. (A)** The expression of selected neural crest genes projected on posterior foxd3+/citrine+ cell UMAP (fig.2 A). **(B)** Relative cell age of posterior foxd3+ cells inferred by scVelo latent time analysis: blue dots correspond to the inferred oldest cells and red – the most recent ones. **(C)** Heatmap displaying *hox* gene expression across cell clusters. **(D)** Heatmap displaying enriched NMp, pro-mesodermal and pro-neural gene expression across NMp-related cell clusters.

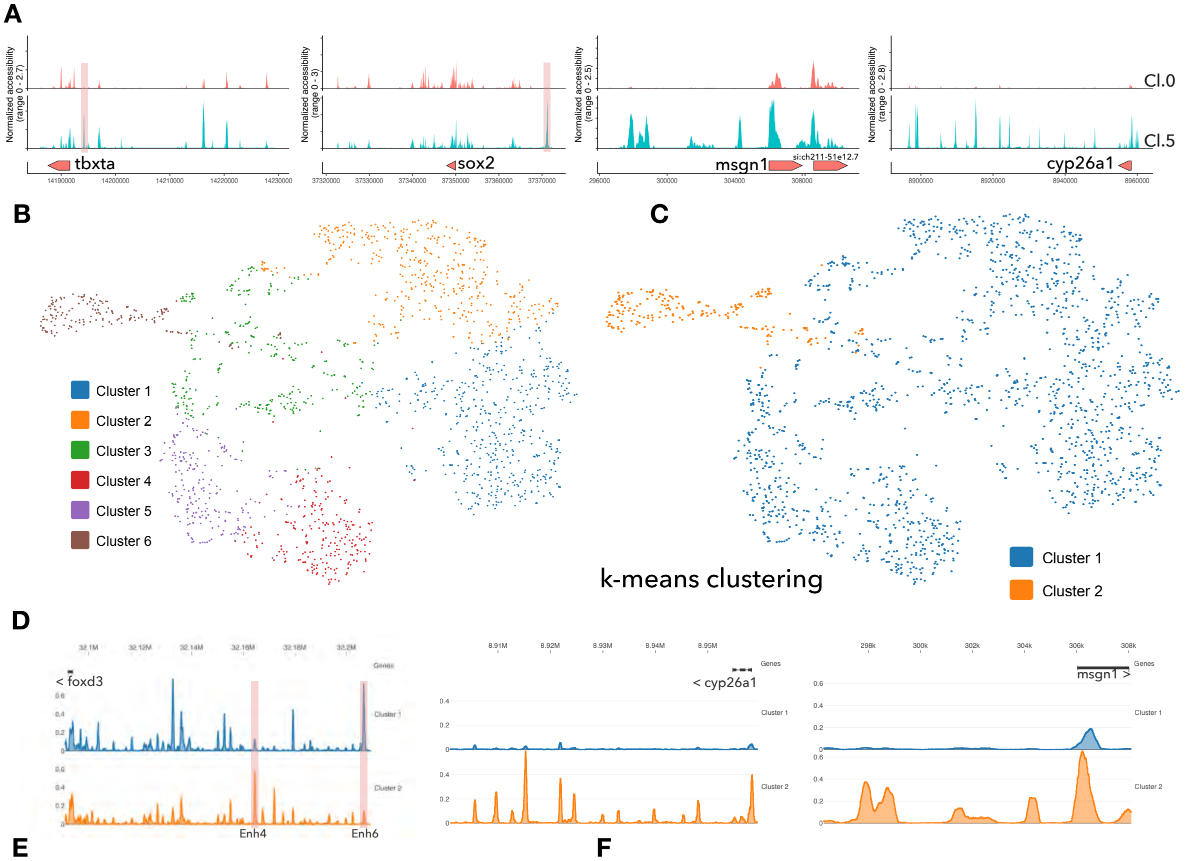


**E**

| Feature name | Cl.1 Av. | Cl.2 Av. | Cl.1 Log2 Fold | P-value |
|---|---|---|---|---|
| sox10 | 8.303081261 | 2.594184264 | 1.681699515 | 3.77E-25 |
| elavl3 | 7.724845229 | 1.96974345 | 1.974031542 | 7.56E-34 |
| AXL | 5.573441869 | 1.643748613 | 1.763488014 | 5.53E-26 |
| tnc | 5.432450458 | 1.721803715 | 1.659777606 | 4.67E-23 |
| foxn3 | 4.335343199 | 1.161643573 | 1.900263222 | 4.01E-28 |
| eya1 | 4.017399004 | 1.235107198 | 1.702262892 | 1.30E-20 |
| sdk2b | 3.911227334 | 0.674947056 | 2.530990325 | 1.09E-37 |
| ets1 | 3.557892582 | 0.987167463 | 1.848978656 | 6.95E-24 |
| col14a1a | 3.467132929 | 1.088179948 | 1.671774649 | 2.19E-21 |
| aatka | 3.428317479 | 0.67035558 | 2.350680323 | 3.10E-32 |
| cavin2b | 3.290180145 | 0.97798451 | 1.749561625 | 3.65E-18 |
| myo10l1 | 3.269059974 | 0.771368064 | 2.08086006 | 2.17E-28 |
| zmiz1a | 2.826107199 | 0.918295315 | 1.620667627 | 1.43E-18 |
| si:ch211-244c8.4 | 2.780441965 | 0.702495916 | 1.981435596 | 1.42E-23 |
| ftr82 | 2.638879738 | 0.849423166 | 1.63369059 | 7.79E-18 |
| lnx1 | 2.560107209 | 0.569343095 | 2.163353339 | 1.53E-26 |
| gsx2 | 2.215905504 | 0.587709001 | 1.909652455 | 2.62E-19 |
| kcnab1a | 2.198781041 | 0.518836853 | 2.076800175 | 5.64E-24 |
| sytl3 | 2.15768233 | 0.33517779 | 2.673022487 | 7.88E-35 |
| igfbp6b | 2.128570743 | 0.528019806 | 2.004902639 | 1.75E-18 |
| comp | 2.008699502 | 0.518836853 | 1.946393418 | 3.83E-19 |

**F**

| Feature name | Cl.1 Av. | Cl.2 Av. | Cl.2 Log2 Fold | P-value |
|---|---|---|---|---|
| hoxc3a | 19.22392209 | 63.95467719 | 1.728422237 | 5.15E-80 |
| mllt3 | 6.221888199 | 28.38450818 | 2.183994706 | 3.56E-100 |
| arid3c | 7.240793744 | 25.80868982 | 1.827994393 | 8.72E-76 |
| hoxc9a | 5.625386074 | 22.61302212 | 2.001490653 | 1.30E-91 |
| hoxc8a | 5.337695096 | 21.85083701 | 2.027762925 | 7.42E-93 |
| cyp26a1 | 3.438592157 | 20.48257699 | 2.568805553 | 7.92E-131 |
| hoxc6b | 5.360527714 | 20.10148444 | 1.901245322 | 4.23E-75 |
| aopep | 5.182433299 | 18.09960065 | 1.798677694 | 1.61E-71 |
| has2 | 3.697742363 | 14.56416369 | 1.972152645 | 7.33E-67 |
| tbx16 | 2.306094342 | 13.9948206 | 2.59570008 | 8.20E-115 |
| dlc | 3.525926918 | 13.91676549 | 1.975206164 | 5.81E-77 |
| cxcl12b | 3.569308891 | 13.80657006 | 1.946101629 | 7.31E-74 |
| plod2 | 2.713656559 | 12.81940259 | 2.234446514 | 1.61E-92 |
| hoxb10a | 3.298742376 | 12.64492648 | 1.933059107 | 4.82E-65 |
| fn1b | 2.751330378 | 12.42912708 | 2.16997159 | 2.79E-90 |
| il17rd | 3.265635081 | 11.87814989 | 1.857394401 | 4.88E-67 |
| hoxb9a | 2.827819646 | 11.63939311 | 2.035745969 | 6.46E-77 |
| aldh1a2 | 2.15140336 | 11.61643573 | 2.427219265 | 3.03E-105 |
| hoxb8a | 2.964815349 | 10.97822049 | 1.883169922 | 1.74E-67 |
| ism1 | 2.390575025 | 10.64763417 | 2.149595202 | 1.39E-71 |
| msgn1 | 2.747905485 | 10.56498759 | 1.937428317 | 5.14E-61 |

S7

### Figure S7

**Teasing out neuromesodermal progenitor (NMp)-related cell specific *cis*-regulatory elements for NMp gene regulatory network building. (A)** Chromatin accessibility tracks across NMp-related genes in scATAC-seq Cl.0 (cranial neural crest (NC)) and Cl.5 (NMps) clusters identified by Signac pipeline. **(B)** Loupe cell browser identified 6 different scATAC-seq clusters based on differential accessibility profiles. **(C)** Loupe cell browser identified the $k$-mean clusters, singles out Cluster 2 as NMp-specific. **(D)** Chromatin accessibility tracks across NMp-related genes in $k$-mean cluster 1 (NC) cluster 2 (NMps), corresponding to the same elements as found in (A). **(E)** Summed accessibility peaks belonging to a nearby gene, enriched in the $k$-mean cluster 1 (NC) over $k$-mean cluster 2 (NMps). **(F)** Summed accessibility peaks belonging to a nearby gene, enriched in the $k$-mean cluster 2 (NMps) over $k$-mean cluster 1 (NC).

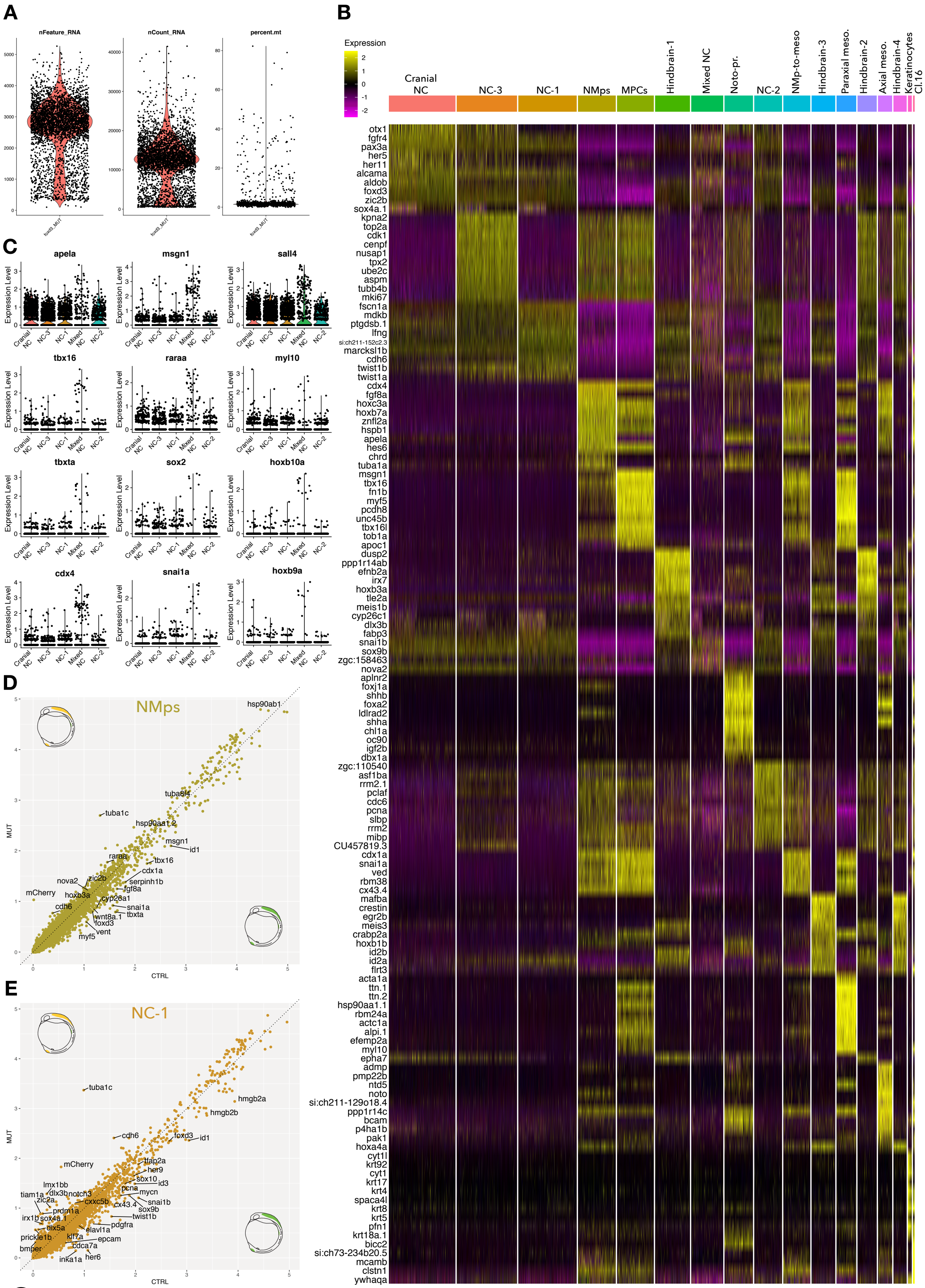
A
nFeature_RNA
nCount_RNA
percent.mt
foxd3_MUT
B
Expression
Cranial NC
NC-3
NC-1
NMps
MPCs
Hindbrain-1
Mixed NC
Noto-pr.
NC-2
NMp-to-meso
Hindbrain-3
Paraxial meso.
Hindbrain-2
Axial meso.
Hindbrain-4
Keratinocytes
Cl.16
otx1
fgfr4
pax3a
her5
her11
alcama
aldob
foxd3
zic2b
sox4a.1
kpna2
top2a
cdk1
cenpf
nusap1
tpx2
ube2c
aspm
tubb4b
mki67
fscn1a
mdkb
ptgdsb.1
lfng
si:ch211-152c2.3
marcksl1b
cdh6
twist1b
twist1a
cdx4
fgf8a
hoxc3a
hoxb7a
znfl2a
hspb1
apela
hes6
chrd
tuba1a
msgn1
tbx16
fn1b
myf5
pcdh8
unc45b
tbx16l
tob1a
apoc1
dusp2
ppp1r14ab
efnb2a
irx7
hoxb3a
tle2a
meis1b
cyp26c1
dlx3b
fabp3
snai1b
sox9b
zgc:158463
nova2
aplnr2
foxj1a
shhb
foxa2
ldlrad2
shha
chl1a
oc90
igf2b
dbx1a
zgc:110540
asf1ba
rrm2.1
pclaf
cdc6
pcna
slbp
rrm2
mibp
CU457819.3
cdx1a
snai1a
ved
rbm38
cx43.4
mafba
crestin
egr2b
meis3
crabp2a
hoxb1b
id2b
id2a
flrt3
acta1a
ttn.1
ttn.2
hsp90aa1.1
rbm24a
actc1a
alpi.1
efemp2a
myl10
epha7
admp
pmp22b
ntd5
noto
si:ch211-129o18.4
ppp1r14c
bcam
p4ha1b
pak1
hoxa4a
cyt1l
krt92
cyt1
krt17
krt4
spaca4l
krt8
krt5
pfn1
krt18a.1
bicc2
si:ch73-234b20.5
mcamb
clstn1
ywhaqa
C
apela
msgn1
sall4
tbx16
raraa
myl10
tbxta
sox2
hoxb10a
cdx4
snai1a
hoxb9a
Expression Level
Cranial NC
NC-3
NC-1
Mixed NC
NC-2
D
NMps
hsp90ab1
tuba8l4
tuba1c
hsp90aa1.2
msgn1
id1
raraa
tbx16
cdx1a
nova2
zic2b
serpinh1b
fgf8a
mCherry
hoxb3a
cyp26a1
cdh6
snai1a
tbxta
wnt8a.1
foxd3
vent
myf5
MUT
CTRL
E
NC-1
tuba1c
hmgb2a
hmgb2b
cdh6
foxd3
id1
mCherry
tfap2a
her9
lmx1bb
id3
sox10
tiam1a
dlx3b
notch3
pcna
mycn
zic2a
cxxc5b
cx43.4
snai1b
prdm1a
sox9b
irx1b
sox4a.1
twist1b
dlx5a
pdgfra
elavl1a
prickle1b
klf7a
epcam
bmper
cdca7a
her6
inka1a
MUT
CTRL

S8

**Figure S8**

**Integrating foxd3+ (citrine+) and foxd3- (citrine+/mCherry+) scRNA-seq datasets.** **(A)** Violin plots of sequenced/unfiltered foxd3-(citrine+/mCherry+) cell quality metrics. **(B)** Top 10 markers enriched in integrated dataset clusters. **(C)** Violin plots depicting neural crest (NC) clusters expressing neuromesodermal progenitor (NMp) related genes. Each dot corresponds to a gene expression value of a single cell. **(D)** Scatter plot displaying average expression of 'NMps' cluster genes by both foxd3+/(citrine+, x-axis) and foxd3- (citrine+/mCherry+, y-axis) cells. **(E)** Scatter plot displaying average expression of 'NC-1' cluster genes by both foxd3+/(citrine+, x-axis) and foxd3- (citrine+/mCherry+, y-axis) cells.

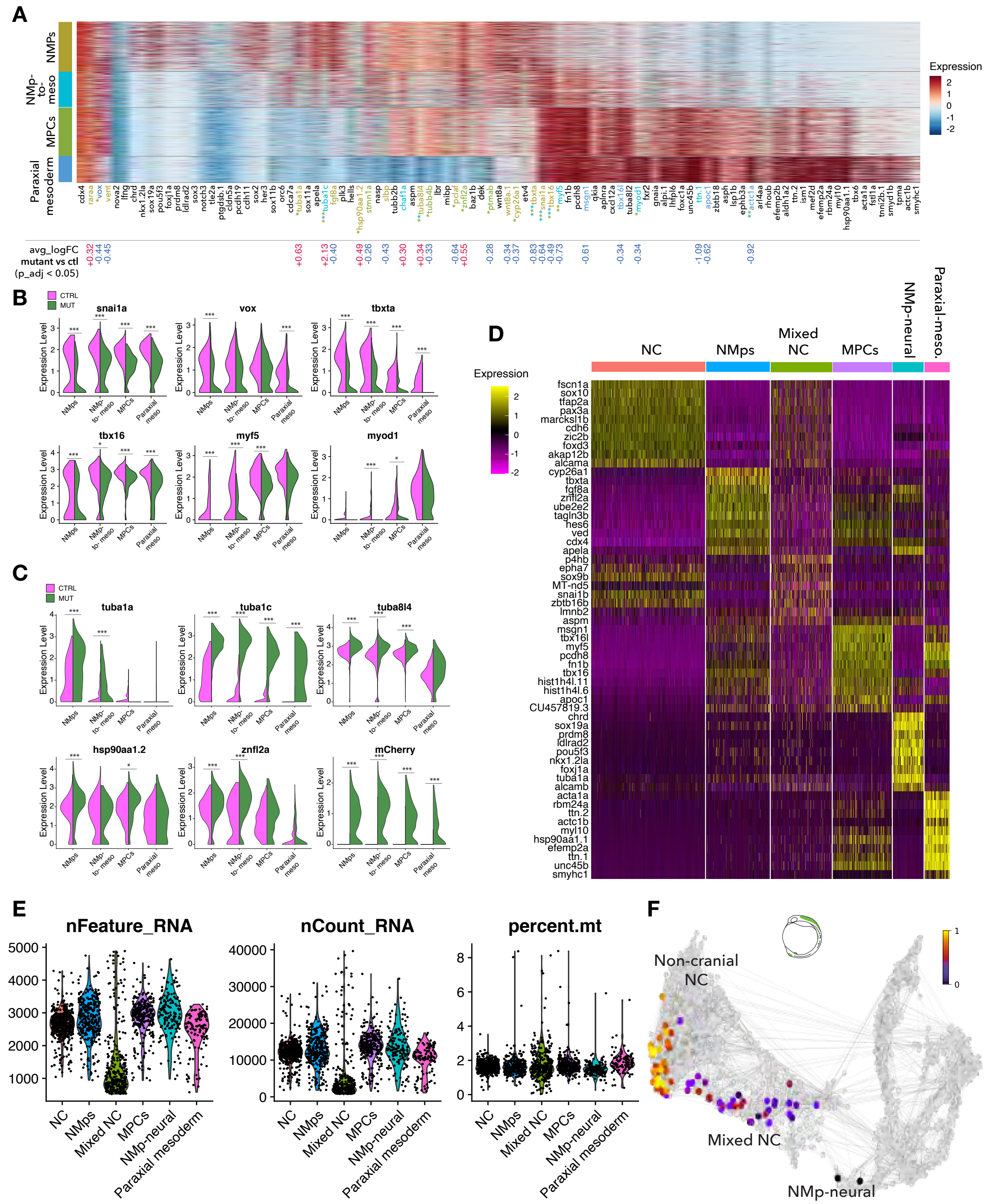
A
NMPs
NMp-to-meso
MPCs
Paraxial mesoderm
Expression
avg_logFC mutant vs ctl (p_adj < 0.05)
B
CTRL
MUT
snai1a
vox
tbxta
tbx16
myf5
myod1
Expression Level
NMps
NMp-to-meso
MPCs
Paraxial meso
C
tuba1a
tuba1c
tuba8l4
hsp90aa1.2
znfl2a
mCherry
D
Expression
NC
NMps
Mixed NC
MPCs
NMp-neural
Paraxial-meso.
fscn1a
sox10
tfap2a
pax3a
marcksl1b
cdh6
zic2b
foxd3
akap12b
alcama
cyp26a1
tbxta
fgf8a
znfl2a
ube2e2
tagln3b
hes6
ved
cdx4
apela
p4hb
epha7
sox9b
MT-nd5
snai1b
zbtb16b
lmnb2
aspm
msgn1
tbx16l
myf5
pcdh8
fn1b
tbx16
hist1h4l.11
hist1h4l.6
apoc1
CU457819.3
chrd
sox19a
prdm8
ldlrad2
pou5f3
nkx1.2la
foxj1a
tuba1a
alcamb
acta1a
rbm24a
ttn.2
actc1b
myl10
hsp90aa1.1
efemp2a
ttn.1
unc45b
smyhc1
E
nFeature_RNA
nCount_RNA
percent.mt
NC
NMps
Mixed NC
MPCs
NMp-neural
Paraxial mesoderm
F
Non-cranial NC
Mixed NC
NMp-neural

S9

**Figure S9**

**Effect on neuromesodermal progenitor (NMp)-related cell development in the absence of foxd3.** **(A)** Heatmap showing NMp and NMp-early derivative gene expression across NMP-related clusters from the foxd3+ (ctl) and foxd3- (mutant) integrated dataset. Average log fold change values of genes that are upregulated in mutant are marked in red and downregulated - in blue. Affected genes are highlighted in cluster-specific colours to specify in which cluster they are significantly upregulated or downregulated. **(B)** Violin plots of differentially expressed pro-medosermal genes between CTRL (foxd3+) or MUT (foxd3-) conditions across NMp-related clusters. Three stars correspond to p ¡ 0.001 and one star to p ¡ 0.05 (Wilcoxon rank sum test between each condition) of downregulated genes in mutant. **(C)** Violin plots of differentially expressed genes between CTRL (foxd3+) or MUT (foxd3-) conditions across NMp-related clusters. Three stars correspond to p ¡ 0.001 and one star to p ¡ 0.05 (Wilcoxon rank sum test between each condition) of upregulated genes in mutant. **(D)** Top 10 markers enriched in posterior foxd3- dataset (fig. 4 E) clusters. **(E)** Violin plots of quality metrics of posterior foxd3- dataset (fig. 4 E) grouped by cluster. **(F)** Predicted descendant/ancestor cells coming from a specified NMp-neural cell in the foxd3+ dataset based on scVelo latent time.

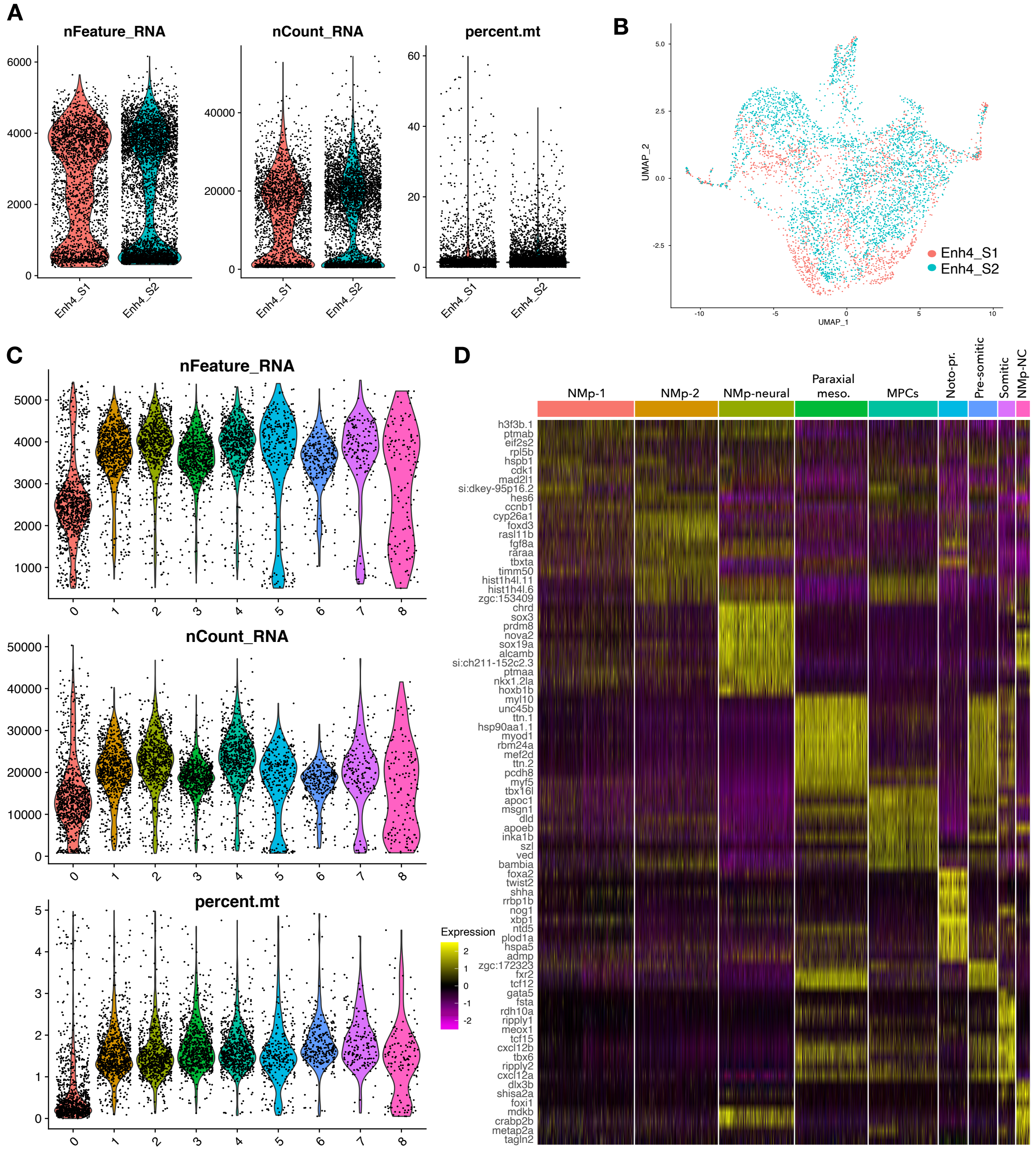
A
nFeature_RNA
nCount_RNA
percent.mt
Enh4_S1
Enh4_S2
B
UMAP_1
UMAP_2
Enh4_S1
Enh4_S2
C
nFeature_RNA
nCount_RNA
percent.mt
D
NMp-1
NMp-2
NMp-neural
Paraxial meso.
MPCs
Noto-pr.
Pre-somitic
Somitic
NMp-NC
Expression
h3f3b.1
ptmab
eif2s2
rpl5b
hspb1
cdk1
mad2l1
si:dkey-95p16.2
hes6
ccnb1
cyp26a1
foxd3
rasl11b
fgf8a
raraa
tbxta
timm50
hist1h4l.11
hist1h4l.6
zgc:153409
chrd
sox3
prdm8
nova2
sox19a
alcamb
si:ch211-152c2.3
ptmaa
nkx1.2la
hoxb1b
myl10
unc45b
ttn.1
hsp90aa1.1
myod1
rbm24a
mef2d
ttn.2
pcdh8
myf5
tbx16l
apoc1
msgn1
dld
apoeb
inka1b
szl
ved
bambia
foxa2
twist2
shha
rrbp1b
nog1
xbp1
ntd5
plod1a
hspa5
admp
zgc:172323
fxr2
tcf12
gata5
fsta
rdh10a
ripply1
meox1
tcf15
cxcl12b
tbx6
ripply2
cxcl12a
dlx3b
shisa2a
foxi1
mdkb
crabp2b
metap2a
tagln2

S10

### Figure S10

**Enh4-EGFP+ scRNA-seq data processing and clustering. (A)** Violin plots of Sample 1 (S1 – 90% epiboly stage) and Sample2 (S2 – 1 somite stage (ss)) sequenced/unfiltered cell quality metrics. **(B)** UMAP embedding for Enh4-EGFP+ cell S1 and S2 scRNA-seq into a single reference. **(C)** Violin plots of integrated S1 and S2 scRNA-seq quality metrics grouped by cluster. **(D)** Top 10 markers enriched in S1+S2 integrated reference clusters; NMp- neuromesodermal progenitor, MPCs – mesodermal progenitor cells, Noto-pr. – notochord progenitors, NC – neural crest.

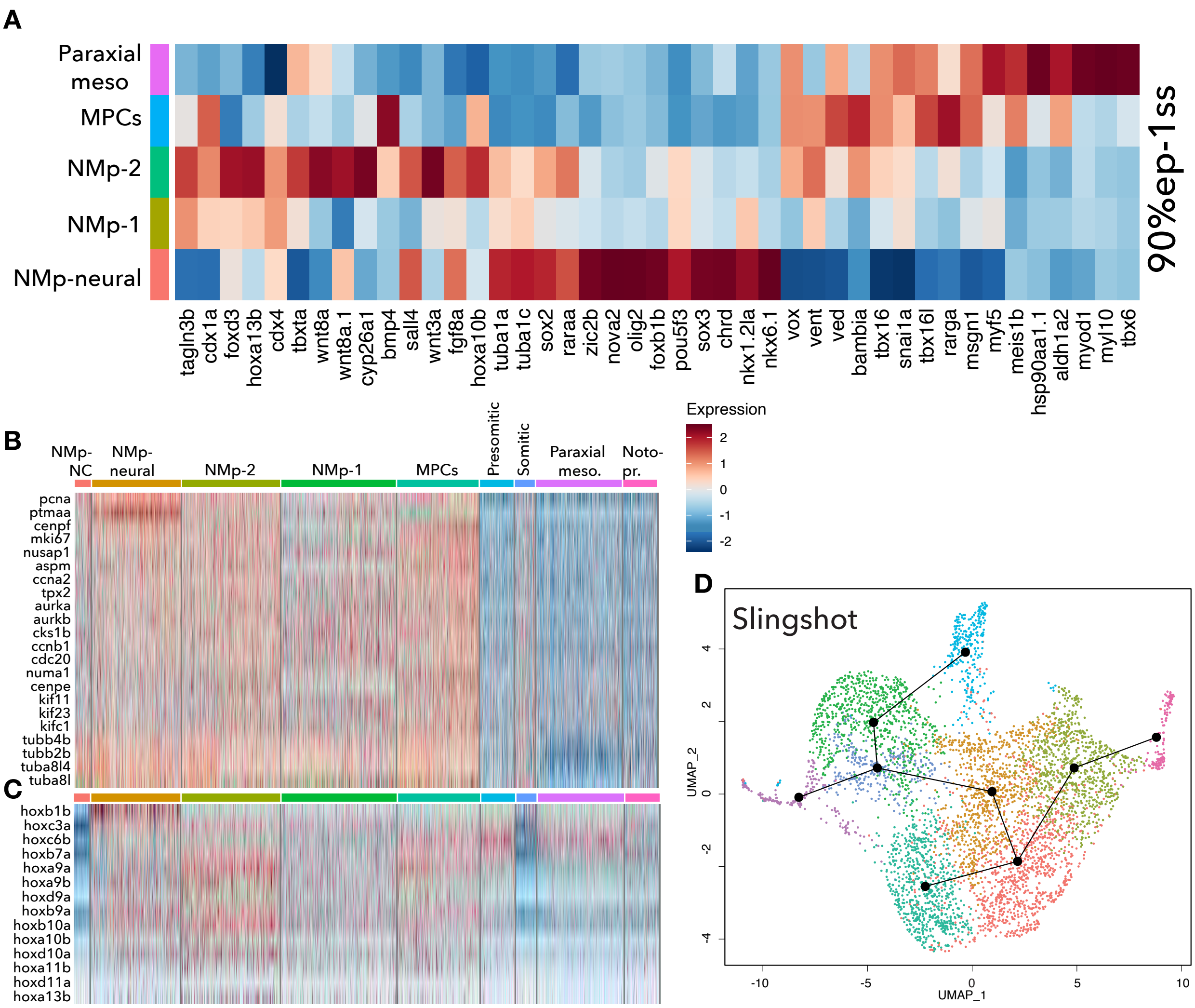


**E**

| Noto-pr. | MPCs | NMP-NC | NMp-1 | NMp-2 | NMp-neural | Paraxial meso | Pre-somitic | Somitic |
|---|---|---|---|---|---|---|---|---|
| raraa | lpar1 | fbxo5 | apela | inka1b | apela | tbx16l | inka1b | ttn.1 |
| qkia | greb1l | rbbp4 | arl5c | arl5c | greb1l | tbx16 | unc45b | pkdcca |
| arl4aa | tbx16 | cavin2b | rbm8a | fbxo5 | fbxo5 | unc45b | tbx16l | large2 |
| arl5c | raraa | arhgap36 | zgc:110158 | FO082779.1 | phkg2 | ttn.1 | arl4aa | FO082779.1 |
| rarga | sulf1 | usp54a | raraa | apela | tle2a | inka1b | pcdh8 | zswim5 |
| zswim5 | greb1 | otx2b | cdx1a | rbbp4 | rbbp4 | lpar1 | thrap3a | inka1b |
| zgc:110158 | gli2a | arhgap27l | fam32a | zswim5 | zswim5 | hsp90aa1.1 | igsf9a | lsp1b |
| chd7 | rreb1b | chd7 | rbbp4 | raraa | arl5c | arl4aa | FO082779.1 | rbbp4 |
| col9a1b | tcf12 | pcna | tle2a | zgc:110158 | cenpf | thrap3a | tbx16 | hsp90aa1.1 |
| letm1 | nrip1a | si:ch211-250c4.4 | ybx1 | fn1b | rarga | rarga | sox5 | slc43a2b |

# S11

### Figure S11

**Early Enh4-EGFP+ cells encompassing neuromesodermal progenitor (NMp), notochord progenitor (noto-pr.) and NMp-neural crest (NC) cells. (A)** Heatmap illustrating selected NMp-related cluster from (fig. 5 A) enriched gene average expression; MPCs – mesodermal progenitor cells. **(B)** Heatmap displaying cell cycling gene expression across all Enh4-EGFP+ cell clusters. **(C)** Heatmap displaying cell *hox* gene expression across all Enh4-EGFP+ cell clusters. **(D)** UMAP embedding illustrating Enh4-EGFP+ integrated dataset with inferred cell lineages (cluster dots connected with lines) by Slingshot. **(E)** scVelo predicted cluster-specific first 10 top-likelihood genes driving cluster/cell transitions based on dynamical modelling.
